# A dual-pathway architecture enables chronic stress to disrupt agency and promote habit formation

**DOI:** 10.1101/2023.10.03.560731

**Authors:** Jacqueline R. Giovanniello, Natalie Paredes, Anna Wiener, Kathia Ramírez-Armenta, Chukwuebuka Oragwam, Hanniel O. Uwadia, Abigail L. Yu, Kayla Lim, Jenna S. Pimenta, Gabriela E. Vilchez, Gift Nnamdi, Alicia Wang, Megha Sehgal, Fernando MCV Reis, Ana C. Sias, Alcino J. Silva, Avishek Adhikari, Melissa Malvaez, Kate M. Wassum

**Author notes:** Correspondence: Kate Wassum Dept. of Psychology, UCLA 1285 Franz Hall, Box 951563, Los Angeles, CA 90095-1563.

## Abstract

Chronic stress can change how we learn and, thus, how we make decisions. Here we investigated the neuronal circuit mechanisms that enable this. Using a multifaceted systems neuroscience approach in male and female mice, we reveal a dual pathway, amygdala-striatal neuronal circuit architecture by which a recent history of chronic stress disrupts the action-outcome learning underlying adaptive agency and promotes the formation of inflexible habits. We found that the basolateral amygdala projection to the dorsomedial striatum is activated by rewarding events to support the action-outcome learning needed for flexible, goal-directed decision making. Chronic stress attenuates this to disrupt action-outcome learning and, therefore, agency. Conversely, the central amygdala projection to the dorsomedial striatum mediates habit formation. Following stress this pathway is progressively recruited to learning to promote the premature formation of inflexible habits. Thus, stress exerts opposing effects on two amygdala-striatal pathways to disrupt agency and promote habit. These data provide neuronal circuit insights into how chronic stress shapes learning and decision making, and help understand how stress can lead to the disrupted decision making and pathological habits that characterize substance use disorders and mental health conditions.

---

When making a decision, we can use what we have learned about our actions and their outcomes to prospectively evaluate the consequences of our potential choices^1–3^. This goal-directed strategy supports our agency. It allows us to choose actions that cause desirable consequences and avoid those that lead to outcomes that are not currently beneficial. This strategy is, thus, highly flexible. But we don’t always think about the consequences of our behavior. Often this is fine. Such habits allow us to efficiently execute routine behaviors based on past success, without forethought of their consequences^4,2,5–7^. The brain balances goal-directed and habitual control to allow behavior to be adaptive when needed, yet efficient when appropriate^8,9^. But disrupted agency and overreliance on habit can cause inadequate consideration of consequences, disrupted decision making, inflexible behavior, and a lower threshold for compulsivity^10–13^. This can contribute to cognitive deficits in numerous diseases, including substance use disorder^14–25^, obsessive-compulsive disorder^26–28^, obesity^12,21,29^, schizophrenia^30–32^, depression^31,33^, anxiety^34^, and autism^35^. Chronic stress tips the balance of behavioral control towards habit^36–60^. Stress can change how we learn and, thus, how we make decisions, attenuating agency and promoting inflexible habits. Because stress is a major predisposing factor for addiction and other psychiatric conditions^61–66^, understanding how stress promotes habit will illuminate one avenue of vulnerability for these conditions. Yet, despite importance for understanding adaptive and maladaptive behavior, little is known of the neuronal circuits that support the learning underlying agency and habits and even less of those that enable stress to potentiate habit formation.

Amygdala-striatal projections are potential candidate pathways by which stress could influence learning and behavioral control strategy. The dorsomedial striatum (DMS) is an evolutionarily conserved hub for the action-outcome learning that supports goal-directed decision making^4,67–75^. Suppression of DMS activity attenuates such agency and promotes inflexible habits^70,76^. The basolateral amygdala (BLA) is also needed for goal-directed behavior^77–79^. It sends a direct excitatory projection to the DMS^80–87^. Little is known of the function of the BLA→DMS pathway, though it is well-positioned to facilitate the action-outcome learning that supports agency. Conversely, the central amygdala (CeA) has been implicated in habit^79^. It sends a direct, likely inhibitory^88,89^, projection to the striatum^80,81,90^ and is, thus, poised to oppose striatal activity. Both the BLA and CeA are highly implicated in stress processing^91–94^. Therefore, here we investigated the function of the BLA→DMS and CeA→DMS pathways in action-outcome and habit learning and asked whether chronic stress acts via these amygdala-striatal pathways to attenuate agency and promote the formation of inflexible habits.

## RESULTS

### Chronic stress disrupts agency and potentiates the formation of inflexible habits

We first designed a behavioral procedure to model stress-potentiated habit formation in male and female mice (Figure 1a). Mice received 14 consecutive days of chronic mild unpredictable stress (“stress”) including daily, pseudorandom, exposure to 2 of 6 stressors: damp bedding (4-16 hrs), tilted cage (4-16 hr), white noise (80 db; 2- 16 hr), continuous illumination during the dark phase (12 hr), physical restraint (2 hr), and footshock (0.7-mA, 1-s, 5 shocks/10 min). This models aspects of the repeated and varied nature of stress experienced by humans, including uncontrollable physical aversive events, disrupted sleep, and poor environmental conditions. Controls received equated handling. Demonstrating efficacy, serum corticosterone was higher (Figure 1b; see Supplemental Table 1 for full statistical reporting) and body weight was lower (Figure 1c) in stressed mice than controls. This procedure was intentionally mild to model low-level, chronic stress. Accordingly, it did not cause major anxiety- or depression-like phenotypes in classic assays of such behavior (Extended Data Figure 1-1). 24 hours following the last stressor, mice were trained to lever press to earn a food-pellet reward. We used 4 sessions of training on a random-ratio schedule of reinforcement in which a variable number of presses (average = 1 - 10, escalated each training session) was required to earn each reward. The tight press-reward relationship of this regime encourages action-outcome learning and, together with the short training duration, the use of such knowledge to support agency and goal-directed decision making^8,95–98^. Mice were food-deprived and bodyweight did not significantly differ between control and stressed mice during training (Supplemental Table 2). Both control and stressed mice similarly acquired the instrumental behavior (Figure 1d). Thus, stress did not cause general learning, motivational, or locomotor impairments. To evaluate behavioral control strategy, we used the gold-standard outcome-specific devaluation test^2–4,7,67,99–102^. Mice were given 90-min, non-contingent access to the food pellet earned during training to induce a sensory-specific satiety rendering that specific food pellet temporarily devalued. Lever pressing was assessed in a 5-min, non-reinforced probe test immediately following the prefeeding. Performance was compared to that following satiation on an alternate food pellet to control for general satiety (Valued state; test order counterbalanced). Both control and stressed mice consumed similar amounts during the prefeed (Supplemental Table 3), indicating that stress did not alter food consumption. Stress also did not affect food pellet discrimination or devaluation efficacy (Supplemental Table 4). If subjects have learned the action-outcome relationship and are using this to support prospective consideration of action consequences for flexible, goal-directed decision making, they will reduce lever pressing when the outcome is devalued. We saw this in control subjects (Figure 1e-f; see also Extended data Figure 1-2 for data on entries into the food-delivery port). Stressed mice were insensitive to devaluation, indicating disrupted agency. Such lack of consideration of action consequences marks inflexible habits^4,7,8,67,99^.

**Figure 1:**
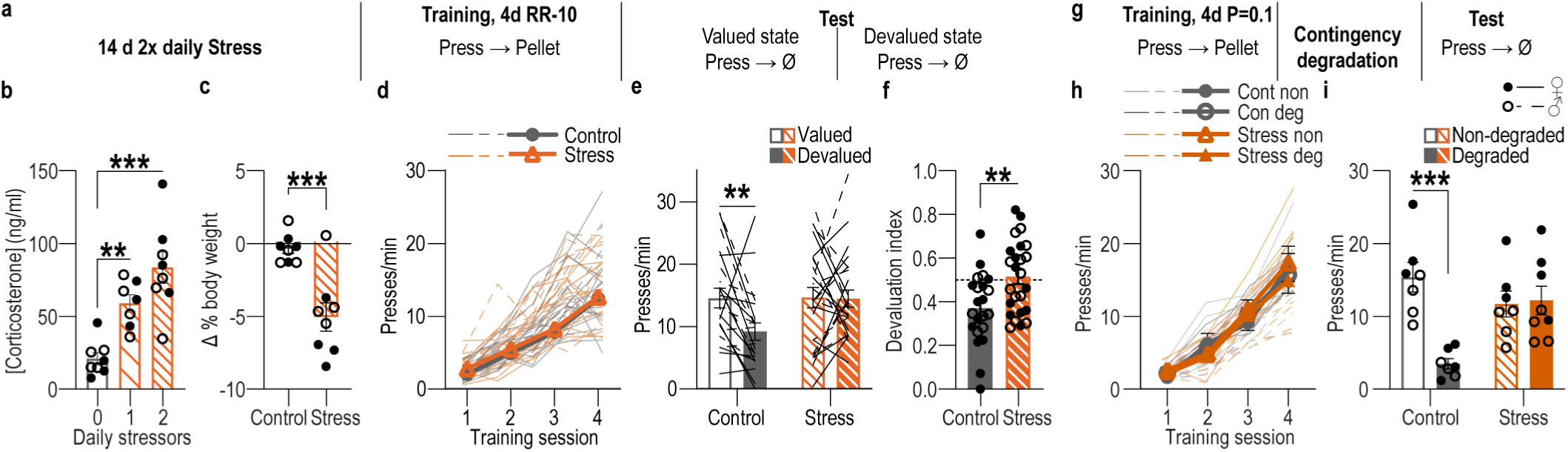
A recent history of chronic stress disrupts action-outcome learning and potentiates habit formation. **(a)** Procedure schematic. Stress, chronic unpredictable mild stress. Lever presses earn food pellet rewards on a random-ratio (RR) reinforcement schedule, prior to lever-pressing probe test in the Valued state, prefed on untrained food-pellet type to control for general satiety, and Devalued state prefed on trained food-pellet type to induce sensory-specific satiety devaluation (order counterbalanced). **(b)** Blood serum corticosterone 24 hr after 14 d of 1 stressor/d, 2 stressors/d, or daily handling (Control). Stress: F_(2, 20)_ = 17.35, *P* < 0.0001. Control N = 8 (4 male), 1x stress N = 7 (3 male), 2x stress N = 8 (4 male). **(c)** Percent change (Δ) in body weight averaged across the first 10 d of stress on ad libitum food. t_14_ = 4.50, *P* = 0.0005. N = 8/group (4 male). **(d)** Press rate across training (beginning with the last day of fixed-ratio 1 training). Training: F_(2.12, 95.32)_ = 168.20, *P* < 0.0001. **(e)** Press rate during the devaluation probe test. Stress x Value: F_(1, 45)_ = 4.43, *P* = 0.04. **(f)** Devaluation index [(Devalued condition presses)/(Valued condition presses + Devalued presses)]. t_(45)_ = 2.99, *P* = 0.005. Control N = 22 (13 male), Stress N = 25 (12 male). **(g)** Procedure schematic. After stress, lever pressing earned pellets with a probability of 0.1. A lever-pressing probe test was conducted following contingency degradation during which presses earned pellets but pellets were also delivered non-contingency absent a press with the same probability or non-degraded control. **(h)** Press rate across training. Training: F_(1.66, 41.39)_ = 211.10, *P* < 0.0001. **(i)** Press rate during the lever-pressing probe test. Stress x Contingency Degradation Group: F_(1, 25)_ = 12.75, *P* = 0.002. Control, Non-degraded N = 7 (3 male), Control, Degraded N = 7 (3 male), Stress Non-degraded N = 7 (3 male) Stress Degraded N = 8 (4 male). Males = closed circles/solid lines, Females = open circles/dashed lines. \*\**P* <0.01, \*\*\**P* < 0.001.

**Figure 2:**
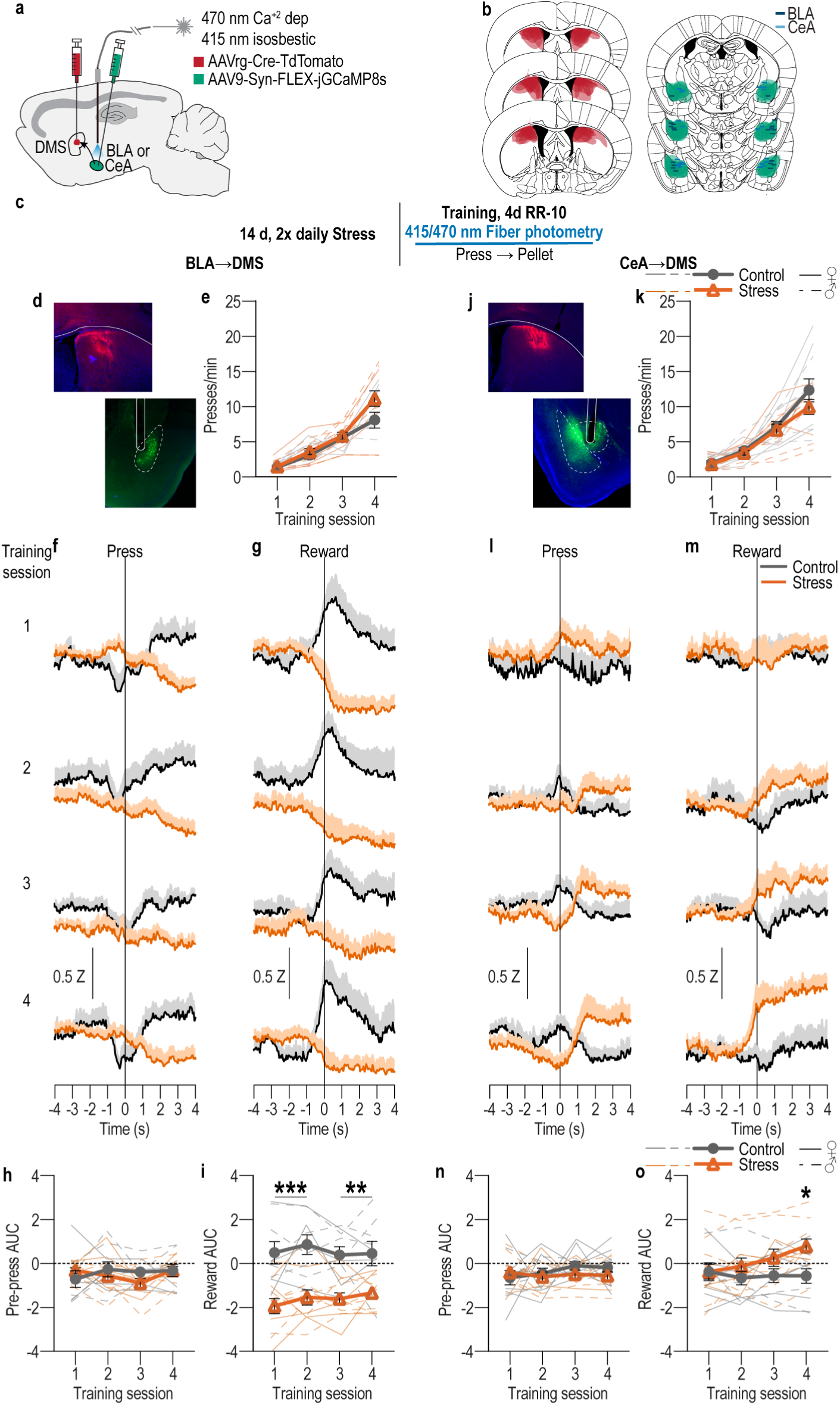
The BLA→DMS pathway is typically activated by rewards during action-outcome learning, chronic stress attenuates this and instead progressively recruits CeA→DMS pathway activity to learning. **(a)** Intersectional approach for fiber photometry calcium imaging of DMS-projecting BLA or CeA neurons. **(b)** Schematic representation of retrograde AAV-cre in DMS and cre-dependent GCaMP8s expression and optical fiber tips in BLA or CeA for all subjects. **(c)** Procedure schematic. Stress, chronic unpredictable stress. Lever presses earned food pellet rewards on a random-ratio (RR) reinforcement schedule. **(d-i)** Fiber photometry recordings of GCaMP8s in BLA→DMS neurons during instrumental lever press – food-pellet reward learning. **(d)** Representative images of retro-cre expression in DMS and immunofluorescent staining of cre-dependent GCaMP8s expression and fiber placement in BLA. **(e)** Press rate across training. Training: F_(1.72, 32.66)_ = 81.40, *P* < 0.0001. **(f-g)** Z-scored Δf/F BLA→DMS GCaMP8s fluorescence changes aligned to bout- initiating presses (f) and reward collection (g) averaged across trials for each subject (shading reflects between-subject s.e.m.) across each training session. **(h-i)** Quantification of area under the BLA→DMS GCaMP8s Z-scored Δf/F curve (AUC) during the 3-s period prior to initiating presses (h; Training: F_(2.49, 47.38)_ = 0.91, *P* = 0.43) or following reward collection (i; Stress: F_(1, 19)_ = 24.13, *P* < 0.0001). Control N = 9 (4 male), Stress N = 12 (5 male). **(j-o)** Fiber photometry recordings of GCaMP8s in CeA→DMS neurons during instrumental lever press – food-pellet reward learning. **(j)** Representative immunofluorescent image of retro-cre expression in DMS and cre-dependent GCaMP8s expression and fiber placement in CeA. **(k)** Press rate across training. Training: F_(1.51, 30.23)_ = 65.61, *P* < 0.0001. **(l-m)** Z-scored Δf/F CeA→DMS GCaMP8s fluorescence changes aligned to bout-initiating presses (l) and reward collection (m) averaged across trials for each subject across each training session. **(n-o)** Quantification of CeA→DMS GCaMP8s Z-scored Δf/F AUC during the 3-s period prior to initiating presses (n; *P* = 0.58; Stress: F_(1, 20)_ = 0.74, *P* = 0.40) or following reward collection (o; Training x Stress: F_(3, 60)_ = 4.51, P = 0.006). Control N = 11 (6 male), Stress N = 11 (4 male). Males = closed circles/solid lines, Females = open circles/dashed lines. \**P* <0.05, \*\**P* <0.01, \*\*\**P* < 0.001.

To provide converging evidence that stress disrupts the action-outcome learning that supports agency, we conducted a second experiment, this time assessing behavioral control strategy using the other gold-standard test: contingency degradation^2–4,100,103^ (Figure 1g). Mice received chronic stress or daily handling control prior to being trained to lever press to earn food-pellet rewards. During training, each press earned reward with a probability that became progressively leaner (1.0 - 0.1). Control and stressed mice, again, similarly acquired the instrumental behavior (Figure 1h). Half the subjects in each group received a 20-min contingency degradation session during which lever pressing continued to earn reward with a probability of 0.1, but reward was also delivered non-contingently with the same probability. Thus, reward was no longer contingent on pressing. The other half received a non-degraded control session in which rewards remained contingent on pressing (see Figure 1-3 for data from the contingency degradation session). Lever pressing was assessed in a 5-min, non-reinforced probe test the next day. If subjects learned the action-outcome contingency and used it to support their agency, their actions should be sensitive to the change in this contingent relationship, such that they will reduce lever pressing when it is no longer needed to earn reward^103^. Controls were sensitive to contingency degradation. Stressed mice were not (Figure 1i). Together these data show that a recent history of chronic stress causes an inability to engage one’s agency and flexibly adapt behavior when its consequence is not currently beneficial or when it is no longer required to earn reward. Thus, chronic stress disrupts action-outcome learning to attenuate agency and, instead, causes the premature formation of inflexible habits.

**Figure 3:**
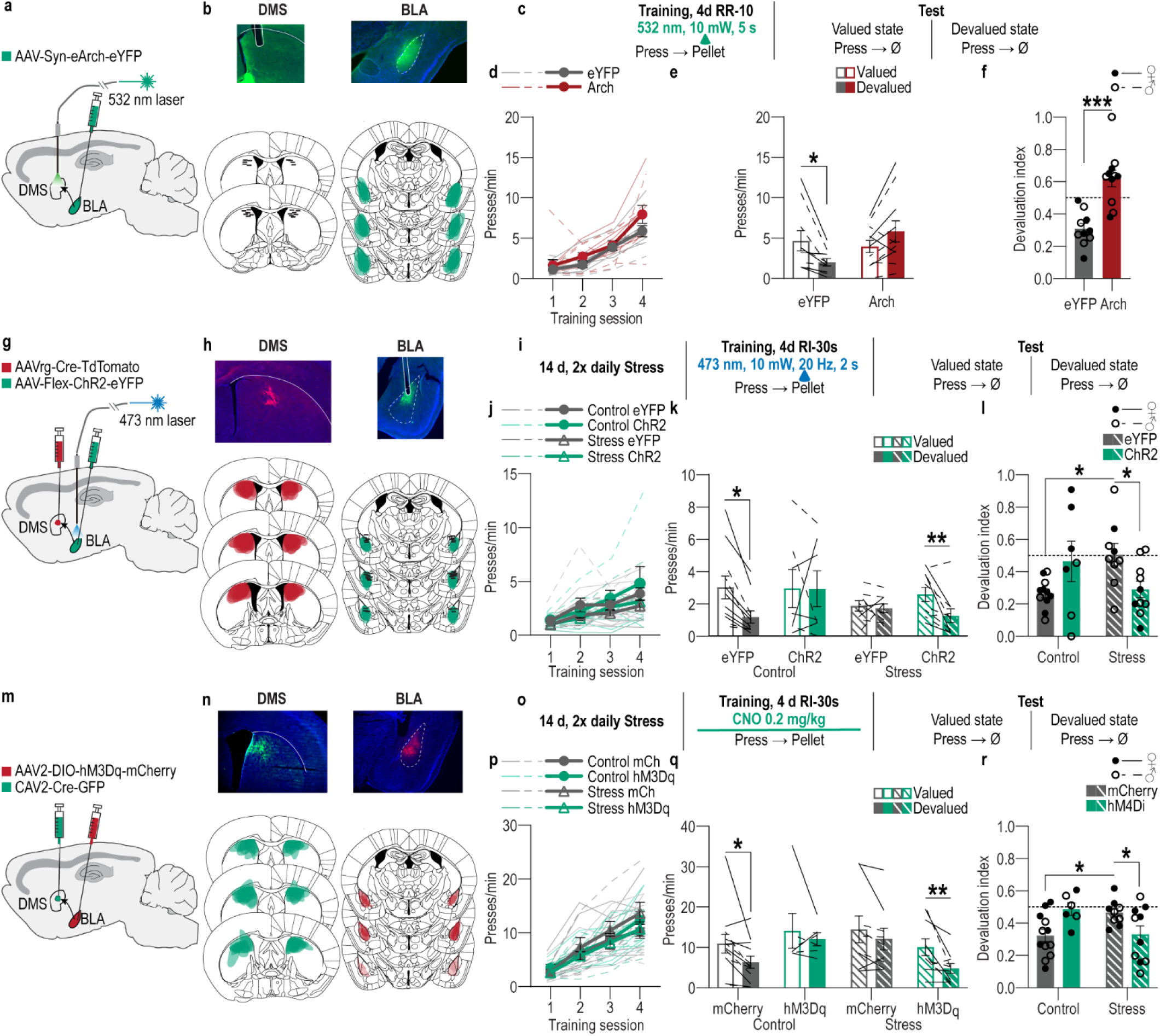
The BLA→DMS pathway mediates action-outcome learning and is suppressed by chronic stress to disrupt agency and promote habit formation. **(a-f)** Optogenetic BLA→DMS inactivation at reward during instrumental learning. **(a)** Approach for optogenetic inhibition of BLA terminals in DMS. **(b)** Top, representative immunofluorescent images of Arch expression in BLA and optical fiber tip in the vicinity of Arch-expressing BLA terminals in DMS. Bottom, schematic representation of Arch expression in BLA and approximate location of optical fiber tips in DMS for all subjects. **(c)** Procedure schematic. Lever presses earned food-pellet rewards on a random-ratio (RR) reinforcement schedule. BLA→DMS projections were optogenetically inhibited at the time of reward during training. Mice were then given a lever-pressing probe test in the Valued state, prefed on untrained food-pellet type and Devalued state prefed on trained food-pellet type to induce sensory-specific satiety devaluation (order counterbalanced). **(d)** Press rate across training. Training: F_(1.70, 32.34)_ = 41.26, *P* < 0.0001. **(e)** Press rate during the devaluation probe test. Stress x Value: F_(1, 19)_ = 14.35, *P* = 0.001. **(f)** Devaluation index [(Devalued condition presses)/(Valued condition presses + Devalued presses)]. t_(19)_ = 5.03, *P* < 0.0001. eYFP N = 10 (5 male), Arch N = 11 (5 male). **(g-l)** Optogenetic BLA→DMS activation at reward during post-stress learning. **(g)** Intersectional approach for optogenetic activation of DMS-projecting BLA neurons. **(h)** Top, representative immunofluorescent images of retro-cre expression in DMS and cre- dependent ChR2 expression in BLA. Bottom, schematic representation of retro-cre in DMS and cre-dependent hM3Dq expression in BLA for all subjects. **(i)** Procedure schematic. Stress, chronic unpredictable stress. After stress, lever presses earned reward on a random-interval (RI) reinforcement schedule. BLA→DMS projections were optogenetically stimulated at the time of reward during training prior to devaluation tests. **(j)** Press rate across training. Training: F_(1.95, 64.18)_ = 30.17, *P* < 0.0001. **(k)** Press rate during the devaluation probe test. Value x Stress x Virus: F_(1, 33)_ = 6.74, *P* = 0.01. *Control groups,* Value x Virus: F_(1, 16)_ = 0.3.13, *P* = 0.10. *Stress groups,* Value x Virus: F_(1, 17)_ = 4.23, *P* = 0.05. **(l)** Devaluation index. Stress x Virus: F_(1, 33)_ = 9.64, *P* = 0.004. Control eYFP N = 11 (7 male), Control ChR2 N = 7 (4 males), Stress eYFP N = 9 (2 male), Stress ChR2 N = 10 Stress (3 male). **(m-r)** Chemogenetic BLA→DMS activation during post-stress learning. **(m)** Intersectional approach for chemogenetic activation of DMS-projecting BLA neurons. **(n)** Top, representative immunofluorescent images of retro-cre expression in DMS and cre-dependent hM3Dq expression in BLA. Bottom, schematic representation of retro-cre in DMS and cre-dependent hM3Dq expression in BLA for all subjects. **(o)** Procedure schematic. After stress, lever presses earned reward on a RI reinforcement schedule. BLA→DMS projections were chemogenetically activated (CNO, clozapine- N-oxide) during training prior to devaluation tests. **(p)** Press rate across training. Training: F_(2.04, 67.36)_ = 73.32, *P* < 0.0001. **(q)** Press rate during the devaluation probe test. Planned comparisons valued v. devalued, Control mCherry: t_(11)_ = 2.76, *P* = 0.01; Control hM3Dq: t_(5)_ = 0.89, *P* = 0.38; Stress mCherry: t_(8)_ = 1.25, *P* = 0.22; Stress hM3Dq: t_(9)_ = 2.9, *P* = 0.007. **(r)** Devaluation index. Stress x Virus: F_(1, 33)_ = 11.60, *P* = 0.002. Control mCherry N = 12 (7 male), Control hm3Dq N = 6 (3 male), Stress mCherry N = 9 (5 male), Stress hM3Dq N = 10 (5 male). Males = closed circles/solid lines, Females = open circles/dashed lines. \**P* < 0.05, \*\**P* < 0.01, \*\*\**P* < 0.001.

### Chronic stress attenuates BLA**→**DMS pathway activity related to action-outcome learning and instead progressively recruits the CeA**→**DMS pathway to learning

We next used both anterograde and retrograde tracing to confirm the existence of direct BLA^80–86^ and CeA^80,81^ projections to dorsal striatum. We found that both BLA and CeA directly project the DMS (Extended data Figure 2-1). We then characterized the activity of these BLA→DMS and CeA→DMS pathways during action-outcome learning and asked whether it is influenced by chronic stress. We used fiber photometry to record fluorescent activity of the genetically encoded calcium indicator GCaMP8s expressed using an intersectional approach in DMS-projecting BLA or CeA neurons (Figure 2a-j). Mice received chronic stress or daily handling control prior to being trained to lever press to earn food-pellet rewards on a random-ratio reinforcement schedule (Figure 2c). Both control and stressed mice similarly acquired the instrumental behavior (Figure 2e, k, see also Extended Data Figure 2-2 for food-port entry data). Fiber photometry (473 nm calcium-dependent, 415 nm isosbestic) recordings were made during each training session. BLA→DMS neurons were robustly activated by earned reward during learning (Figure 2f-i). Thus, the BLA→DMS pathway is active when subjects are able to link the rewarding consequence to their actions, thus forming the action-outcome knowledge that supports agency. This activity was absent in stressed mice (Figure 2g, i). Chronic stress attenuates the BLA→DMS activity associated with action-outcome learning. Conversely, CeA→DMS neurons were not robustly active during this form of instrumental learning in control subjects, indicating CeA→DMS projection activity is not associated with action-outcome learning. Stress caused the CeA→DMS pathway to be progressively engaged around earned reward experience with training (Figure 2l-o). The CeA→DMS response to earned reward was long lasting, taking approximately 30 seconds to return to baseline after reward collection (Extended Data Figure 2-3). Thus, a recent history of chronic stress causes the CeA→DMS pathway to be recruited to instrumental learning. We detected similar patterns in response to unpredicted rewards in both pathways (Extended Data Figure 2-4). Both BLA→DMS and CeA→DMS projections were acutely activated by unpredicted aversive events (footshock; Extended Data Figure 2-4), indicating that neither BLA→DMS nor CeA→DMS bulk activity is valence-specific. These aversive responses were not altered by stress (Extended Data Figure 2-4), providing a positive control for our ability to detect signal in all groups. Chronic stress did, however, reduce post-shock fear-related BLA→DMS activity, consistent with its effects on reward signals in this pathway. Chronic stress did not alter baseline spontaneous calcium activity in either pathway, indicating it does not generally increase or decrease excitability in these pathways (Extended Data Figure 2-5). Together these data indicate that a recent history of chronic stress oppositely modulates BLA→DMS and CeA→DMS pathway activity. BLA→DMS projections are normally activated by rewarding events, but stress prevents this learning-related activity and, instead, causes the CeA→DMS pathway to be progressively recruited during learning.

### The BLA**→**DMS pathway mediates action-outcome learning and is suppressed by chronic stress to disrupt agency and promote habit formation

#### BLA→DMS projections are activated by rewards to support action-outcome learning for flexible, goal-directed decision making

Reward experience is an opportunity to link the reward to the action that earned it, forming the action-outcome knowledge that supports agency. BLA→DMS projections are activated by earned rewards. So, we reasoned that this activity might be critical for action-outcome learning. If this is true, then inhibiting reward- evoked BLA→DMS activity should suppress action-outcome learning and, thereby, disrupt flexible goal-directed decision making. We tested this by optogenetically inhibiting BLA→DMS projection activity at the time of earned reward during instrumental learning. We expressed the inhibitory opsin archaerhodopsin (Arch) or fluorophore control in the BLA and implanted optical fibers in the DMS in the vicinity of Arch-expressing BLA axons and terminals (Figure 3a-b). Mice were trained to lever press to earn food-pellet rewards on a random-ratio schedule of reinforcement. We optically (532 nm, 10 mW, 5 s) inhibited BLA terminals in the DMS during each earned reward (Figure 3c). BLA→DMS inhibition did not affect acquisition of the instrumental behavior (Figure 3d; see also Extended Data Figure 3-1 for food-port entry data). Training was followed by a set of outcome-specific devaluation tests, as above. No manipulation was given at test to allow us to isolate BLA→DMS function in action-outcome learning rather than the expression of such learning during decision making. Controls were sensitive to outcome devaluation, indicating action-outcome learning for goal-directed decision making. Inhibition of BLA→DMS projections during learning caused subsequent insensitivity to outcome devaluation (Figure 3e-f). BLA→DMS inhibition was not inherently rewarding or aversive (Extended Data Figure 3-2). Thus, BLA→DMS projections are normally activated by rewarding events to enable the action-outcome learning that supports agency.

#### Stress-induced suppression of BLA→DMS projections disrupts action-outcome learning and enables premature habit formation

Since, BLA→DMS projections are critical for action-outcome learning, we next reasoned that the stress-induced suppression of BLA→DMS activity might disrupt such learning. We tested this by asking whether activating BLA→DMS projections during learning, to counter the effects of stress, is sufficient to restore action-outcome learning and, thus, goal-directed decision making in stressed mice. We did this in two ways. Because chronic stress abolishes reward-evoked BLA→DMS activity during learning, we first used optogenetics to stimulate BLA→DMS projections at the time of earned reward during learning following chronic stress. Using an intersectional approach (Figure 3g), we expressed the excitatory opsin Channelrhodopsin 2 (ChR2) or fluorophore control in DMS-projecting BLA neurons (Figure 3h). Following chronic stress or daily handling control, mice were trained to lever press to earn food-pellet rewards. We used a random interval schedule of reinforcement in which a variable (average 30-s) period of time had to elapse after an earned reward before a press would earn another reward. Limited training on this regime allows action-outcome learning for goal-directed decision making^99^. But the looser action-outcome relationship is more permissive for habits than a ratio reinforcement schedule^96,97,102,104^, thereby making it more difficult to neurobiologically prevent stress-potentiated habit and the results more robust if such an effect were to occur. We optically (473 nm, 10 mW, 20Hz, 2 s) stimulated DMS-projecting BLA neurons during collection of each earned reward (Figure 3i). Neither stress nor BLA→DMS stimulation significantly altered acquisition of the instrumental behavior (Figure 3j). Training was followed by the outcome-devaluation test, conducted without manipulation. Whereas controls were sensitive to subsequent outcome devaluation, indicating action-outcome learning and flexible goal-directed decision making, stressed mice were insensitive to devaluation, indicating premature habit formation (Figure 3-l). Optogenetic activation of BLA→DMS projections during learning restored normal action-outcome learning enabling agency, as evidenced by sensitivity to devaluation, in stressed mice (Figure 3k-l). Thus, activation of BLA→DMS projections during reward learning is sufficient to overcome the effect of prior chronic stress and restore action-outcome learning to enable agency for flexible, goal-directed decision making.

To provide converging evidence, we conducted a second experiment in which we activated the BLA→DMS pathway during post-stress learning using chemogenetics. Using an intersectional approach (Figure 3m), we expressed the excitatory designer receptor human M3 muscarinic receptor (hM3Dq) or fluorophore control in DMS-projecting BLA neurons (Figure 3n). Following chronic stress or daily handling control, mice were trained to lever press to earn food-pellet rewards on a random-interval reinforcement schedule (Figure 3o). Prior to each instrumental training session, mice received the hM3Dq ligand clozapine-N-oxide (CNO; 0.2 mg/kg^105–108^ i.p.) to activate BLA→DMS projections. Neither stress nor chemogenetic BLA→DMS activation altered instrumental acquisition (Figure 3p). Mice then received devaluation tests. Whereas controls were sensitive to outcome devaluation, stressed mice were, again, insensitive to devaluation, indicating premature habit formation (Figure 3q-r). Chemogenetic activation of BLA→DMS projections during learning replicated the effect of optogenetic activation, restoring action-outcome learning to enable goal-directed decision making, as evidenced by sensitivity to devaluation, in stressed mice (Figure 3q-r). Neither optogenetic nor chemogenetic activation of BLA→DMS projections significantly impacted learning in subjects without a history of chronic stress. Though behavior was variable in these groups with some marginal evidence of an influence on action-outcome learning, perhaps due to disruption of neurotypical activity. Together these data reveal that BLA→DMS projections are activated by rewards to enable the action-outcome learning that supports flexible, goal-directed decision making and chronic stress attenuates this to disrupt such agency and promote premature habit formation.

### The CeA**→**DMS pathway mediates routine habit formation and is recruited by chronic stress to promote premature habit

#### CeA→DMS projections mediate the formation of routine habits

The CeA is necessary for habit^79^. This function may be achieved, at least in part, via its direct inhibitory projection to DMS. It is, therefore, perhaps not surprising that the CeA→DMS pathway is not typically active during action-outcome learning. Rather the CeA is activated by rewards following overtraining^109^. Therefore, we reasoned that the CeA→DMS pathway might mediate the natural habit formation that occurs for routine behaviors. To test this, we asked whether CeA→DMS projection activity is necessary for habit formation by optogenetically inhibiting CeA→DMS projections at the time of earned reward during learning and overtraining. We expressed the inhibitory opsin Arch or fluorophore control in the CeA and implanted optical fibers in the DMS in the vicinity of Arch-expressing CeA axons and terminals (Figure 4a-c). Mice were trained to lever press to earn food-pellet rewards on a random interval schedule of reinforcement and were overtrained to promote natural habit formation (Figure 4c). We optically (532 nm, 10 mW, 5 s) inhibited CeA terminals in the DMS during each earned reward (Figure 4c). Training was followed by the outcome-specific devaluation test. No manipulation was given on test to allow us to isolate CeA→DMS function in habit learning rather habit expression. Optogenetic CeA→DMS inhibition did not alter acquisition of the instrumental behavior (Figure 4d; see also Extended Data Figure 4-1 for food-port entry data). It did, however, prevent habit formation. Controls showed evidence that they formed routine habits, insensitivity to devaluation. But mice for which we inhibited the CeA→DMS pathway during overtraining continued to show flexible goal-directed decision making, sensitivity to devaluation (Figure 4e-f). Thus, the CeA→DMS pathway mediates the natural habit formation that occurs with repeated practice of an instrumental routine.

**Figure 4:**
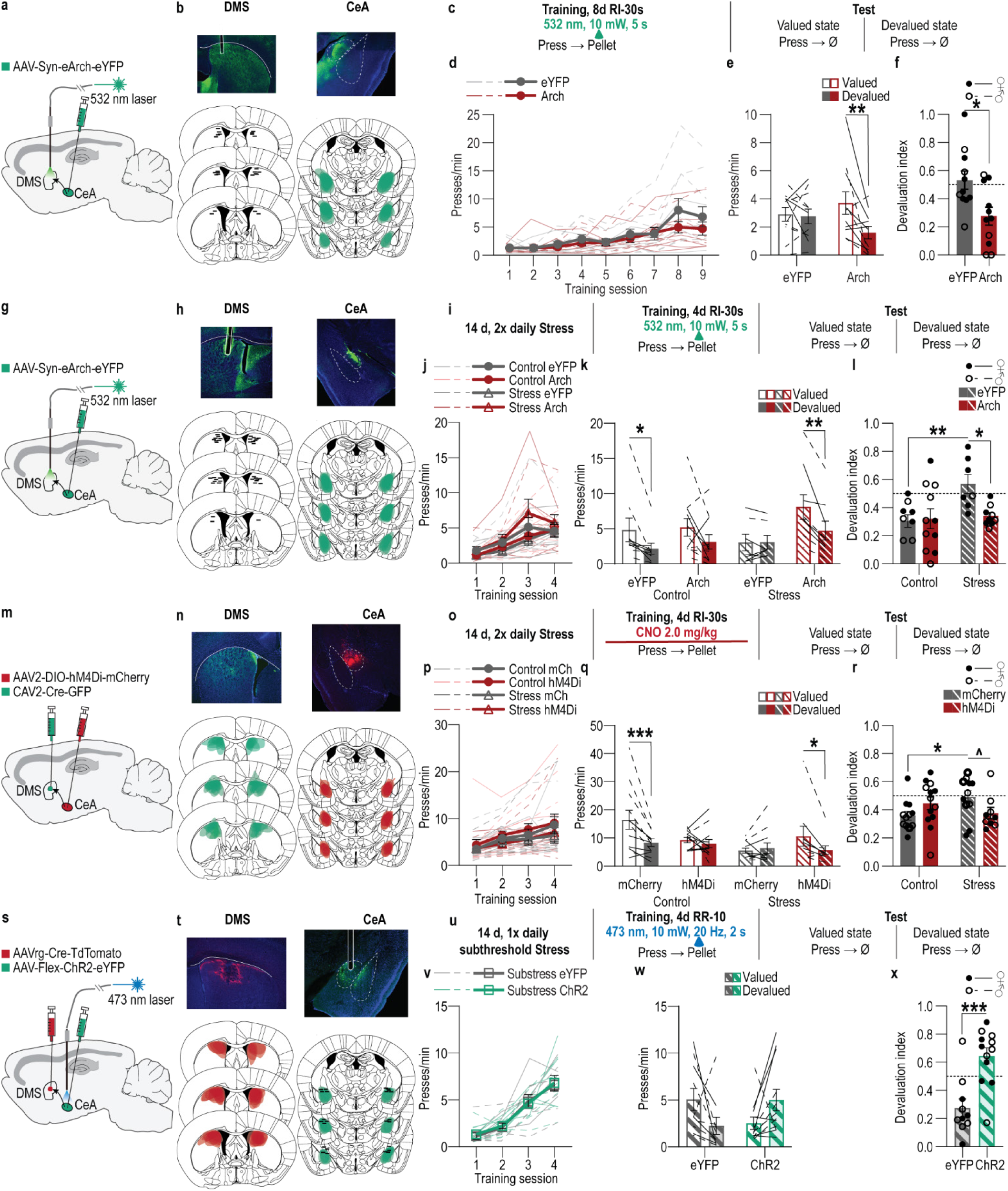
The CeA→DMS pathway mediates habit formation and is recruited by chronic stress to promote premature habits. **(a-f)** Optogenetic inactivation of CeA→DMS projections at reward during natural habit formation. **(a)** Approach for optogenetic inhibition of CeA terminals in DMS. **(b)** Top, representative immunofluorescent images of Arch expression in CeA and optical fiber tip in the vicinity of Arch-expressing CeA terminals in the DMS. Bottom, schematic representation of Arch expression in BLA and approximate location of optical fiber tips in DMS for all subjects. **(c)** Procedure schematic. Lever presses earned food pellet rewards on a random-interval (RI) reinforcement schedule. Mice were overtrained to promote habit formation. CeA→DMS projections were optogenetically inactivated during each training session prior to lever-pressing probe tests in the Valued state, prefed on untrained food-pellet type, and Devalued state prefed on trained food-pellet type to induce sensory-specific satiety devaluation (order counterbalanced). **(d)** Press rate across training. Training: F_(1.46, 29.09)_ = 15.69, *P* = 0.0001. **(e)** Press rate during the devaluation probe test. Virus x Value: F_(1, 20)_ = 4.72, *P* = 0.04. **(f)** Devaluation index [(Devalued condition presses)/(Valued condition presses + Devalued presses)]. t_(20)_ = 2.80, *P* = 0.01. eYFP N = 11 (3 male), Arch N = 11 (7 male). **(g-l)** Optogenetic CeA→DMS inactivation at reward during post-stress learning. **(g)** Approach for optogenetic inhibition of CeA terminals in DMS. **(h)** Top, representative immunofluorescent images of Arch expression in CeA and optical fiber tip in the vicinity of Arch-expressing CeA terminals in the DMS. Bottom, schematic representation of Arch expression in BLA and approximate location of optical fiber tips in DMS for all subjects. **(i)** Procedure schematic. Stress, 2x daily chronic unpredictable stress. Lever presses earned food-pellet rewards on a RI reinforcement schedule. CeA→DMS projections were optogenetically inactivated at reward during training, prior to devaluation tests. **(j)** Press rate across training. Training: F_(2.15, 68.91)_ = 31.05, *P* < 0.0001. **(k)** Press rate during the devaluation probe test. Value x Stress x Virus: F_(1, 32)_ = 4.14, *P* = 0.05. *Control groups,* Value x Virus: F_(1, 18)_ = 0.15, *P* = 0.70. *Stress groups,* Value x Virus: F_(1, 14)_ = 12.88, *P* = 0.003. **(l)** Devaluation index. Stress x Virus: F_(1, 32)_ = 4.47, *P* = 0.04. Control eYFP N = 9 (5 male), Control Arch N = 11 (4 male), Stress eYFP N = 7 (6 male), Stress Arch N = 9 (5 male). **(m-r)** Chemogenetic CeA→DMS inhibition during post-stress learning. **(m)** Intersectional approach for chemogenetic inhibition of DMS-projecting CeA neurons. **(n)** Top, representative immunofluorescent images of retro-cre expression in DMS. Bottom, cre-dependent hM4Di expression in CeA and schematic representation of retro-cre in DMS and cre-dependent hM4Di expression in CeA for all subjects. **(o)** Procedure schematic. After stress, lever presses earned food-pellet rewards on a RI reinforcement schedule. CeA→DMS projections were chemogenetically inactivated (CNO, clozapine-N-oxide) during training, prior to devaluation tests. **(p)** Press rate across training. Training: F_(1.54, 63.31)_ = 21.12, *P* < 0.0001. **(q)** Press rate during the devaluation probe test. Planned comparisons valued v. devalued, Control mCherry: t_(11)_ = 4.59, *P* < 0.0001; Control hM3Dq: t_(12)_ = 0.73, *P* = 0.46; Stress mCherry: t_(10)_ = 0.47, *P* = 0.64; Stress hM3Dq: t_(8)_ = 2.41, *P* = 0.02. **(r)** Devaluation index. Stress x Virus: F_(1, 41)_ = 5.99, *P* = 0.02. Control mCherry N = 12 (5 male), Control hM4Di N = 13 (8 male), Stress mCherry N = 11 (5 male), Stress hM4Di N = 9 (4 male). **(s-x)** Optogenetic CeA→DMS stimulation at reward during learning following subthreshold chronic stress. **(s)** Intersectional approach for optogenetic stimulation of DMS-projecting CeA neurons. **(t)** Top, representative images of retro- cre expression in DMS and immunofluorescent staining of cre-dependent ChR2 expression in CeA. Bottom, schematic representation of retro-cre in DMS and cre-dependent ChR2 expression in CeA for all subjects. **(u)** Procedure schematic. Subthresold stress, 1x daily chronic unpredictable stress. Lever presses earned food pellet rewards on a random-ratio (RR) reinforcement schedule. CeA→DMS projections were optogenetically activated at the time of reward during training, prior to devaluation tests. **(v)** Press rate across training. Training: F_(2.30, 45.90)_ = 71.93, *P* < 0.0001. **(w)** Press rate during the devaluation probe test. Virus x Value: F_(1, 20)_ = 7.40, *P* = 0.01. **(x)** Devaluation index. t_(20)_ = 3.29, *P* = 0.0004. eYFP N = 10 (4 male), ChR2 N = 12 (6 male). Males = closed circles/solid lines, Females = open circles/dashed lines. ns = not significant. ^*P* = 0.069, \**P* < 0.05, \*\**P* < 0.01, \*\*\**P* <0.001.

#### Stress-induced recruitment of CeA→DMS projections mediates premature habit formation

Given that the CeA→DMS pathway mediates habit formation, we next reasoned that the stress-induced recruitment of this pathway to learning may enable stress to promote premature habit formation. If this is true, then preventing the stress-induced increase in CeA→DMS activity during learning should prevent premature habit formation and restore action-outcome learning and, therefore, agency. We tested this in two ways. Because chronic stress engages the CeA→DMS pathway at reward experience during learning, we first optogenetically inhibited CeA→DMS projections at the time of earned reward during learning following stress. We expressed the inhibitory opsin Arch or fluorophore control in the CeA and implanted optical fibers in the DMS (Figure 4g-h). Following chronic stress or daily handling control, mice were trained to lever press to earn food-pellet rewards and we optically (532 nm, 10 mW, 5 s) inhibited CeA terminals in the DMS during each earned reward (Figure 4i). We used a random interval schedule of reinforcement to increase the robustness of the results. Neither stress nor CeA→DMS inhibition altered acquisition of the instrumental behavior (Figure 4j). Training was followed by outcome-specific devaluation tests, conducted without manipulation. At test, we again found evidence of goal-directed decision making, sensitivity to devaluation, in control subjects and potentiated habit formation, insensitivity to devaluation, in stressed subjects (Figure 4k-l). Optogenetic inhibition of CeA→DMS activity at reward during learning restored action-outcome learning to enable goal-directed decision making in stressed mice, as evidenced by sensitivity to devaluation (Figure k-l). Thus, stress-induced activation of CeA→DMS projections during reward learning is necessary to promote premature habit formation.

To provide converging evidence, we conducted a second experiment in which we chemogenetically inhibited CeA→DMS projections during learning following stress. We used an intersectional approach (Figure 4m) to express the inhibitory designer receptor human M4 muscarinic receptor (hM4Di) or a fluorophore control in DMS-projecting CeA neurons (Figure 4m-n). Following chronic stress or daily handling control, mice were trained to lever press to earn food-pellet rewards on a random interval reinforcement schedule (Figure 4o). Prior to each training session, mice received the hM4Di ligand CNO (2.0 mg/kg^110–112^ i.p.) to inactivate CeA→DMS projections. Neither stress nor chemogenetic CeA→DMS inactivation altered acquisition of the instrumental behavior (Figure 4p). Chemogenetic inhibition of CeA→DMS projections during learning replicated the effects of optogenetic inhibition, restoring action-outcome learning to enable goal-directed decision making, sensitivity to devaluation, in stressed mice (Figure 4q-r). Neither optogenetic nor chemogenetic CeA→DMS inhibition significantly impacted learning or behavioral control strategy in subjects without a history of chronic stress. Together, these data indicate that chronic stress engages CeA→DMS projections during subsequent reward learning experience to promote the premature formation of inflexible habits.

#### CeA→DMS projection activity is sufficient to promote premature habit formation following subthreshold chronic stress

We next asked whether CeA→DMS pathway activity at reward during learning is sufficient to promote habit formation. We used an intersectional approach (Figure 4s) to express the excitatory opsin ChR2 or a fluorophore control in DMS-projecting CeA neurons and implanted optic fibers above the CeA (Figure 4t). We first optically (473 nm, 10 mW, 20 Hz, 25-ms pulse width, 2 s) stimulated CeA→DMS neurons during earned reward during instrumental learning on a random-ratio schedule of reinforcement in mice without a history of chronic stress. This neither affected acquisition of the lever-press behavior, nor the action-outcome learning needed to support flexible, goal-directed decision making during the devaluation test (Extended Data Figure 4-2). Thus, activation of the CeA→DMS pathway during reward learning experience is not alone sufficient to disrupt action-outcome learning or promote habit formation.

We next reasoned that activation of CeA→DMS projections might be sufficient to tip the balance of behavioral control towards habit in the context of a very mild stress experience. We repeated the experiment this time in mice with a history of once daily stress for 14 consecutive days (Figure 4s-u). Again, neither CeA→DMS activation nor stress altered acquisition of the instrumental behavior (Figure 4v). The less frequent chronic stress was itself insufficient to cause premature habit formation. Mice were sensitive to devaluation, indicating preserved action-outcome learning and agency (Figure 4w-x). But activation of CeA→DMS projections at reward during learning was sufficient to cause premature habit formation, as evidenced by greater insensitivity to devaluation in subjects that received stimulation relative to those that did not (Figure 4w-x). Thus, activation of CeA→DMS projections during learning is sufficient to amplify the effects of prior subthreshold chronic stress to promote habit formation. CeA→DMS stimulation was not inherently rewarding or aversive in either control or stressed subjects (Figure 4-3). Together, these data indicate that chronic stress recruits the CeA→DMS pathway to subsequent learning to promote the premature formation of inflexible habits.

## DISCUSSION

These data reveal a dual pathway neuronal circuit architecture by which a recent history of chronic stress shapes learning to disrupt adaptive agency and promote inflexible habits. Both the BLA and CeA send direct projections to the DMS. The BLA→DMS pathway is activated by rewarding events to support the action-outcome learning needed for flexible, goal-directed decision making. Chronic stress attenuates this activity to disrupt action-outcome learning and, therefore, agency. Conversely, the CeA→DMS pathway mediates habit formation. Stress recruits this pathway to learning to promote the premature formation of inflexible habits. Thus, chronic stress disrupts agency and promotes habit formation by altering the balance of BLA and CeA input to the DMS.

Here we provide a model for the function of amygdala-striatal projections. Whereas the BLA→DMS pathway mediates action-outcome learning to support agency, the CeA→DMS pathway mediates the formation of routine habits. BLA→DMS pathway function in action-outcome learning is consistent with evidence that BLA lesion or BLA/DMS disconnection disrupts goal-directed behavior^77–79 82^. We implicate direct BLA→DMS projections and show that this pathway is activated by rewarding events to link those rewards to the actions that earned them to enable the prospective consideration of action consequences needed for flexible decision making. These data do not accord with evidence that BLA→DMS ablation does not disrupt action-outcome learning^113^. Such ablations may allow compensatory mechanisms, that are not possible with temporally-specific manipulation. Unlike the BLA→DMS pathway, the CeA→DMS pathway is not typically activated during action-outcome learning. Rather CeA→DMS projections mediate the natural habit formation that occurs with repeated practice of routine. This is consistent with evidence that CeA neurons are activated by rewards with overtraining^109^, that CeA lesion disrupts habit^79^, and that CeA→DMS projections oppose flexible adjustment of behavior when an action is no longer rewarded^90^. Unlike valence-processing models of amygdala function^114,115^, our data indicate that BLA and CeA projections to DMS are unlikely to convey simple positive or negative valence, but rather differentially shape the content of learning. The data support a parallel model^116^ whereby, via distinct outputs to the DMS, the amygdala actively gates the nature of learning to regulate the balance of behavioral control strategies. An important question opened by this model is how different reward learning experiences, schedules of reinforcement, and training regimes recruit activity in these pathways and how this intersects with stress and other life experiences.

Stressful life events can disrupt one’s agency and promote the formation of inflexible, potentially maladaptive, habits. Indeed, after chronic stress, people become less able to adapt their behavior when its outcome has been devalued^36,38,40,43,46,47^. Using two independent tests, we provide evidence in male and female mice that a recent history of chronic stress disrupts the action-outcome knowledge needed for agency and instead causes the premature formation of inflexible habits. We find that chronic stress disrupts action-outcome learning and promotes habit formation by flipping the balance of BLA and CeA inputs to the DMS.

Chronic stress attenuates reward-learning-related activity in the BLA→DMS pathway to disrupt action-outcome learning and agency and instead recruits activity in the CeA→DMS pathway to promote the formation of inflexible habits. That agency could be rescued by manipulations to oppose these stress effects during only the learning phase indicates that stress influences behavioral control by shaping learning. The stress-induced attenuation of BLA→DMS activity was surprising because the BLA is, generally, hyperactive following chronic stress^117–124^ (c.f.^125^). This may suggest that the effects of stress on the BLA neurons depends on their projection target. Elevated CeA→DMS activity following stress is consistent with evidence that stress increases CeA activity^126–131^. Whereas, stress attenuated BLA→DMS activity throughout learning, the CeA→DMS pathway was progressively recruited across training in stressed subjects. This could indicate that stress-induced CeA→DMS engagement requires repeated reward learning or reinforcement opportunity. It could also suggest the CeA→DMS pathway is engaged to compensate for the stress-induced attenuation of the BLA→DMS pathway activity needed for action-outcome learning. Indeed, the transition of behavioral control to habit systems requires a shift of behavioral control from BLA to CeA^132^. Such speculations require further evaluation of amygdala-striatal activity using dual-pathway recordings and manipulations. Interestingly, activation of the CeA→DMS pathway was not sufficient itself to promote habit formation. CeA→DMS activation did, however, tip the balance towards habit following a subthreshold mild chronic stress experience. Thus, stress may prime the CeA→DMS pathway to be recruited during subsequent learning. CeA→DMS activation may work along with a confluence of disruptions, likely to the BLA→DMS pathway, but also to cortical inputs to DMS^60,104,133^ to promote habit formation. The CeA can also work indirectly, likely via the midbrain^134–137^, with the dorsolateral striatum to regulate habit formation^79,132^. Thus, the CeA may promote habit through both direct and indirect pathways to the striatum. Although evidence from the terminal optogenetic inhibition experiments confirm involvement of direct amygdala projections to DMS, both pathways may collateralize and such collaterals may, too, be involved in learning and affected by stress.

The discoveries here open the door to many important future questions. One is the mechanisms through which prior chronic stress affects amygdala-striatal activity. That chronic stress occurred before training and did not alter spontaneous activity in either pathway, suggests that it may lay down neuroplastic changes in these pathways that become influential during subsequent learning opportunities. How such changes occur is a big and important question for future research. They likely involve a combination of stress action in the amygdala, perhaps via canonical stress systems such as corticotropin releasing hormone^138^ and/or kappa/dynorphin^139^, and stress action at regions upstream to the amygdala. Epigenetic mechanisms may also be involved^140^. An equally substantial next question is how these pathways influence downstream DMS activity. Indeed, DMS neuronal activity, especially plasticity in dopamine D1 receptor-expressing neurons^141^, is critical for the action-outcome learning that supports goal-directed decision making^70–72,104^ and when suppressed promotes inflexible habits^70,76^. A reasonable speculation is that the excitatory BLA→DMS pathway promotes downstream learning-related activity in DMS to support action-outcome learning and that the inhibitory CeA→DMS pathway dampens such activity to encourage habit formation. Amygdala-striatal inputs may coordinate in this regard with corticostriatal inputs known to be important for supporting action-outcome learning^22,60,142^ and susceptible to chronic stress^60^. Both amygdala subregions and DMS participate in drug-seeking^132,143–148^ and active-avoidance behavior^149^. The central amygdala is particularly implicated in compulsive drug seeking and drug seeking after extended use, dependence and withdrawal, or stress^132,147,150,151^. Thus, more broadly, our results indicate chronic stress could oppositely modulate BLA→DMS and CeA→DMS pathways to promote maladaptive drug-seeking and/or avoidance habits. Towards this end, whether individual differences in the balance of BLA→DMS and CeA→DMS activity confer resilience or susceptibility to stress-potentiated habit formation is an important future question.

Adaptive decision making often requires understanding your agency in a situation. Knowing that your actions can produce desirable or undesirable consequences and using this to make thoughtful, deliberate, goal-directed decisions. Chronic stress can disrupt agency and promote inflexible, habitual control over behavior. We found that stress does this with a one-two punch to the brain. Chronic stress dials down the BLA→DMS pathway activity needed to learn the association between an action and its consequence to enable flexible, well-informed decisions. It also dials up activity in the CeA→DMS pathway, causing the formation of rigid, inflexible habits. These data provide neuronal circuit insights into how chronic stress shapes how we learn and, thus, how we decide. This helps understand how stress can lead to the disrupted decision making and pathological habits that characterize substance use disorders and mental illness.

## AUTHOR CONTRIBUTIONS

JRG and KMW conceptualized and designed the experiments, interpreted the data, and wrote the paper. JRG executed all experiments and analyzed the data. NP, AW, CO, HOU, ALY, JSP, GN, GEV assisted with experiments. MS assisted with rabies tracing experiments with resources from AJS. FMCVR assisted with RTPP experiments, with advice and resources from AA. ACS wrote initial code for photometry analysis. KRA analyzed spontaneous event frequency and amplitude on photometry data. MM contributed to the conceptualization and design of the experiments, initially optimized the instrumental conditioning procedures and data analysis, and provided important contributions to the interpretation of the data.

## ACKNOWLEDGEMENTS

This research was supported by NIH R01DA046679 (KMW), NIH R01DA035443 (KMW), NIH T32DA024635 (JRG), NIH F32DA056201 (JRG), A.P. Giannini Fellowship (JRG), NIH K99MH135177 (JRG), NIH TL4GM118977 (NP), NIH R01MH119089 (AA), and the Staglin Center for Behavior and Brain Sciences. UCLA Behavioral Testing Core provided space and behavioral testing equipment for the open field, elevated plus maze and light-dark emergence test.

## COMPETING FINANCIAL INTERESTS

The authors have no biomedical financial interests or potential conflicts of interest to declare.

## METHODS

See Supplemental Table 5 for key reagents.

### Subjects

Male and female wildtype C57/Bl6J mice (Jackson Laboratories, Bar Harbor, ME) aged 9-12 weeks old at the time of surgery served as subjects. Rabies tracing was conducted with *Drd1a-Cre* and *Adora2A-Cre* transgenic mice bred in house and aged 8-16 weeks at the time of surgery. Mice were housed in a temperature (68-79 °F) and humidity (30-70%) regulated vivarium on 12:12 hour reverse dark/light cycle (lights off at 7 AM). Behavioral experiments were performed during the dark phase. Mice were group housed in same-sex groups of 3-4 mice/cage prior to onset of behavioral experiments and subsequently single-housed for the remainder of the experiment to facilitate food deprivation and preserve implants. Unless noted below, mice were provided with food (standard rodent chow, Lab Diet, St. Louis, MO) and water *ad libitum* in the home cage. Mice were handled for 3-5 days prior to the start of behavioral training for each experiment. All procedures were conducted in accordance with the NIH Guide for the Care and Use of Laboratory Animals and were approved by the UCLA Institutional Animal Care and Use Committee.

### Surgery

Mice were anesthetized with isoflurane (3% induction, 1% maintenance), and positioned in a digital stereotaxic frame (Kopf, Tujunga, CA). Subcutaneous Rimadyl (Carprofen; 5 mg/kg; Zoetis, Parsippany, NJ) was given pre- operatively for analgesia and anti-inflammatory purposes. Small cranial holes (1–2 mm^2^) were drilled, through which virus or fluorescent tracers were delivered via a guide cannula (DMS: 28 ga, BLA/CeA: 33 ga), PlasticsOne, Roanoke, VA) connected to a 1-mL syringe (Hamilton Company, Reno, NV) by intramedic polyethylene tubing (BD; Franklin Lakes, NJ) and controlled by a syringe pump (Harvard Apparatus, Holliston, MA). Coordinates (from Bregma) were determined by mouse brain reference atlas^152^ and were as follows: CeA, AP -1.2, ML ±2.8, DV -4.6 mm; BLA, AP -1.5, ML ±3.2, DV -5.0 mm; DMS, AP +0.2, ML ±1.8, DV -2.65 mm.

Virus or tracers were infused at a rate of 0.1 µL/min and cannulae were left in place for at least 10 min post-injection. For injection-only surgeries, the skin was re-closed with Vetbond tissue adhesive (3M, Saint Paul, MN). For surgeries requiring fiber-optic cannulae, fibers were placed 0.3 mm above the target region for optogenetic experiments and at the infusion site for fiber photometry experiments, secured to the skull using RelyX Unicem Universal Self-Adhesive Resin (3M) and a head cap was created using C&B Metabond quick adhesive cement system (Parkell Inc., Brentwood, NY), followed by opaque dental cement (Lang Dental Manufacturing, Wheeling, IL). After surgery, mice were kept on a heating pad maintained at 35 °C for 1 hour and then single-housed in a new homecage for recovery and monitoring. Mice received chow containing the antibiotic TMS for 7 days following surgery to prevent infection, after which they were returned to standard rodent chow. Specific surgical details for each experiment are described below. In all cases, surgery occurred prior to the onset of stress and/or behavioral training.

### Chronic mild unpredictable stress

The chronic mild unpredictable stress (“stress”) procedure was modified from^60,153–156^. Mice assigned to the stress group were exposed to 2 stressors/day (foot shock, physical restraint, tilted cage, white noise, continuous illumination, or damp bedding) for 14 days in a pseudorandomized manner at variable time onset and for varying durations between 2 and 16 hours. Control subjects received equated daily handling in the vivarium by the same experimenter administering the stress. Stress was administered in a separate, enclosed laboratory space distinct from both the vivarium and behavioral testing rooms. Stressed mice had home-cage nesting material removed for the duration of the stress exposure^157^. Mice were transported to the stress space in individual 16-oz clear polyethylene containers and on a dedicated transport cart and placed into individual cages in the stress space. Stress efficacy was assessed by daily body weight measurements^158^. Sub-threshold stress exposure was identical to stress except mice received only 1 stressor/day.

#### Stressors

##### Footshock

Subjects were placed in the conditioning chamber for 2 min to acclimate and then exposed to 5, 2-3 s, 0.7-mA footshocks with a variable intertrial interval averaging 60 sec (30-90 sec range). The footshock chamber had a similar grid floor to the behavioral testing chambers (described below) but was otherwise distinct in wall shape (round), pattern (monochrome polka dot), lack of bedding, scent (75% ethanol), and lighting (off). The chambers also lacked food ports and levers. Chambers were cleaned with 75% ethanol between subjects.

##### Physical Restraint

Subjects were immobilized in modified 50-mL polypropylene conical tubes with 4 air holes per side, 1 at the top, and 1 in the cap for the tail (10 total). Mice were scruffed and placed inside the conical tube for 2 hours in their stress cage.

##### Tilted cage

Stress cages were placed on chocks to tilt each cage at approximately a 45-degree angle for 6 - 16 hours.

##### White noise

100-db white noise was played in the stress space for all stress mice for a duration of 6 - 16 hours.

##### Continuous illumination

Overhead lights were turned on during the dark phase of the light cycle (7PM – 7AM).

##### Damp bedding

∼200 mL of water was mixed with the stress cage corncob bedding. Mice were placed in their stress cage with this damp bedding for 6 - 16 hours. Mice were returned to a new home cage with clean, dry bedding afterwards.

#### Corticosterone ELISA

*N* = 11 male and *N* = 12 female mice were used for corticosterone measurements of blood serum after exposure to 0, 1, or 2 stressors per day for 14 day. Measurements were taken 24 hours after the final stress exposure. Mice were decapitated and trunk blood was collected in 1.7-mL sample tubes on ice. Tubes were centrifuged at 2000 g for 10 min at 4 °C. Clear supernatant was collected and placed in new 1.7-mL sample tubes and frozen at -20 °C. Samples were diluted 1:40 in sample dilution buffer. Serum corticosterone levels were assessed using a Corticosterone ELISA kit as directed (Enzo Biosciences; Farmingdale, NY) and quantified on a microplate reader (Molecular Devices, San Jose, CA).

### Behavioral procedures

#### Instrumental conditioning and tests

Instrumental conditioning procedures were adapted from our prior work^99^.

##### Apparatus

Training took place in Med Associates wide mouse conditioning chambers (East Fairfield, VT) housed within sound- and light-attenuating boxes. Each chamber had metal grid floors and contained a retractable lever to the left of a recessed food-delivery port (magazine) on the front wall. A photobeam entry detector was positioned at the entry to the food port. Each chamber was equipped with 2 pellet dispensers to deliver either 20-mg grain or chocolate-flavored purified pellets (Bio-Serv, Frenchtown, NJ) into the food port. A fan mounted to the outer chamber provided ventilation and external noise reduction. A 3-watt, 24-volt house light mounted on the top of the back wall opposite the food port provided illumination. To monitor subject behavior, monochrome digital cameras (Med Associates) were positioned over top of the conditioning chambers. For optogenetic manipulations, chambers were outfitted with an Intensity Division Fiberoptic Rotary Joint (Doric Lenses, Quebec, QC, Canada) connecting the output fiber optic patch cords to a 473-nm or 593-nm laser (Dragon Lasers, ChangChun, JiLin, China) positioned outside of the chamber.

##### Food deprivation

3 - 5 days prior to the start of behavioral training, mice were food-deprived to maintain 85%- 90% of their free-feeding body weight. Mice were given 1.5 - 3.0 g of their home chow at the same time daily at least 2 hours after training sessions. For experiments involving stress, food deprivation began during the last 3 days of the stress procedure. Owing to food deprivation, body weights did not differ between groups at the start or end of training (see Supplemental Table 2).

##### Outcome pre-exposure

To familiarize subjects with the food pellet that would become the instrumental outcome, mice were given 1 session of outcome pre-exposure. Mice were placed in a clean, empty cage and allowed to consume 20 - 30 of the food pellets from a metal cup. If any pellets remained, they were placed in the home cage overnight for consumption.

##### Magazine conditioning

Mice received 1 session of training in the operant chamber to learn where to receive the food pellets (20-mg grain or chocolate-purified pellets). Mice received 20 - 30 non-contingent pellet deliveries from the food port with a fixed 60-s intertrial interval.

##### Instrumental conditioning

Mice received 4 sessions (1 session/day consecutively), minimum, of instrumental conditioning in which lever presses earned delivery of a single food pellet. Earned pellet type (grain or chocolate) was counterbalanced across subjects within each group of each experiment. Each session began with the illumination of the house light and extension of the lever, and ended with the retraction of the lever and turning off of the house light. Sessions ended after the total available outcomes (20 or 30, as noted for each experiment below) had been earned or a maximum time limit (20 or 30 min, as noted below) had been reached. In all cases, training began on a fixed-ratio 1 schedule (FR-1), in which each action was reinforced with one food pellet outcome. Once mice completed 2 sessions in which they achieved 80% performance criteria (earned 80% of the max outcomes), the reinforcement schedule was escalated to either random interval (RI) or random ratio (RR) as described for each experiment below. For the RI protocol, mice received 1 session on an RI-15 s schedule then 2 - 3 sessions on the final RI-30 s schedule (variable average 15-s or 30-s interval must elapse following a reinforcer for another press to be reinforced). Subjects on the RR protocol received 1 session each of RR-2, RR- 5, and RR-10 schedule of reinforcement (variable press requirement average of 2, 5, or 10 presses to earn the food pellet). For the overtraining protocol, mice received 8 total training sessions, 1 on an RI-15 s schedule then 7 sessions on the final RI-30 s schedule.

For subjects in the contingency degradation experiment, following FR-1 training, they received 2 days of training in which each press was reinforced with a probability of 0.2 and a final session in which each press earned reward with a probability of 0.1.

##### Alternate outcome exposures

To equate exposure of the non-trained pellet, all mice were given non-contingent access to same number of the alternate food pellets (e.g., chocolate pellets if grain pellets served as the training outcome) as the earned pellet type in a different context (clear plexiglass cage) a minimum of 2 hours before or after (alternated daily) each RI or RR instrumental training session.

##### Sensory-specific satiety outcome devaluation test

Testing began 24 hr after the final instrumental conditioning session. Mice were given 1 - 1.5 hours access to either 4 g of the food pellets previously earned by lever pressing (Devalued condition) or 4 g of the non-trained pellets to control for general satiety (Valued condition). The remaining pellets were weighed following prefeeding to measure total consumption. Consumption did not significantly differ between the Devalued v. Valued conditions for any experiment (Supplemental Table 3). Immediately after this prefeeding, lever pressing was assessed during a brief, 5-min, non-reinforced probe test. Following the probe test, mice were given a 10-min consumption choice test with simultaneous access to 1 g of both pellet types to ensure rejection of the devalued outcome. In all cases, mice consumed less of the prefed pellet than non-prefed pellet, indicating successful sensory-specific satiety devaluation (Supplemental Table 4). 24 hr after the first devaluation test, mice received 1 session of instrumental retraining on the final reinforcement schedule (RI30 or RR10), followed the next day by a second devaluation test in which they were prefed the opposite food pellet. Thus, each mouse was tested in both the Valued and Devalued conditions, with test order counterbalanced across subjects within each group for each experiment.

##### Contingency degradation test

24 hr after the final instrumental conditioning session, mice received a 20-min contingency degradation session during which lever pressing continued to earn reward with a probability of 0.1, but reward was also delivered freely with the same probability even if mice did not press (i.e., non-contingent). Thus, lever pressing was no longer necessary to earn reward. This session was identical for non-degraded controls, except they did not receive non-contingent rewards. 24 hr following the contingency degradation session, the effects of this contingency change were assessed in a brief, 5-min non-reinforced probe test.

#### Real-time place preference/avoidance test

Procedure was conducted as described previously^159^. Mice were habituated to a 2-sided opaque plexiglass chamber (20 × 42 × 27 cm) for 10 min, during which their baseline preference for the left or right side of the chamber was measured. During the first 10-min test session, one side of the chamber was assigned to the light-delivery side (counterbalanced across subjects within each group). Mice were placed in the non-stimulation side to start the experiment. Light (Dragon Laser; Changchun, China) was delivered upon entry into the light-paired side and continued until the subject exited that side (optical stimulation: 473 nm, 5-ms pulse width, 20 Hz, ∼8- 10 mW at fiber tip; optical inhibition: 593 nm, continuous, ∼8-10 mW). Mice then received a second test, identical to the first, in which the opposite side of the chamber served as the light-paired side. Sessions were video-recorded using a CCD camera. This camera interfaced with Biobserve software (Biobserve GmbH, Germany) and a Pulse Pal (Sanworks, Rochester, NY), to track subject position in real time and trigger laser delivery. The apparatus was cleaned with 75% ethanol after each session. Distance traveled, movement velocity, and time spent in each chamber was generated by Biobserve software post-session. Time spent in laser-paired chamber was compared between groups to assess preference or aversion of laser delivery.

#### Open-field test

Procedure was conducted as described previously^159^. Mice were placed in an opaque plexiglass arena (34 × 34 × 34 cm) for a single 10-min session. Sessions were video recorded using a CCD camera interfaced with Anymaze (Stoelting Co., Wood Dale, IL) software, which was used to track subject position in real time. Center region was defined as the innermost third of the floor area. Brightness above the OFT was ∼70 lux. The apparatus was cleaned with 75% ethanol after each subject. Distance traveled, movement velocity, and time spent in either center or surrounding outer area was generated by Bioserve software and compared between groups.

#### Light/dark emergence test

The dark side of a 2-chamber apparatus was made of black opaque plexiglass and completely enclosed except for a small entry through the middle divider. The light side was made of white opaque plexiglass and was open to the light above. Brightness in the light chamber was ∼70 lux. Mice were placed in the open portion of the apparatus to initiate a 10-min session. Each session was video recorded using a CCD camera, which interfaced with Anymaze software to track subject location. The apparatus was cleaned with 75% ethanol after each session. Distance traveled, movement velocity, and time spent in the light chamber was generated by Biobserve software and compared between groups.

#### Elevated plus maze

Procedure was conducted as described previously^160^. The dimensions of the elevated plus maze (EPM) arms were 30 cm x 7 cm, and the height of the closed arm walls was 20 cm. The maze was 65 cm elevated from the floor and was placed in the center of the behavior room away from other stimuli. Brightness above the EPM was ∼70 lux. For the 10-min EPM test, mice were placed in the center of EPM facing a closed arm. Each session was video recorded using a CCD camera, which interfaced with Anymaze software to track subject location in real time. The apparatus was cleaned with 75% ethanol after each session. Distance traveled, movement velocity, and time spent in the center, open arms, or closed arms was generated by Biobserve software and compared between groups.

#### Sucrose-preference test

Mice first received habituation to 2 standard home-cage water bottles filled with water in the home cage for 16 hours. Subsequently, one water bottle was replaced with a bottle of 10% sucrose. Bottles were left in place for 24 hours and weighed before and after placement. Bottle positions were switched for another 24-hour period and subsequently weighed again. Amount of sucrose and water consumed, as well as a ratio of the two, during the 48-hour period was compared between groups.

#### Progressive ratio test

Mice were trained on the instrumental training protocol above to a reinforcement schedule of RR-10. They were then given a progressive ratio test in which the number of lever presses required to receive a pellet increased by 4 with each reinforcer delivered (e.g., 1, 5, 9, 13, 17, 21 etc.). The session ended after >5-min break in pressing or maximum duration of 4 hr. Session duration, rewards delivered, total presses, and the break point (last completed press requirement) were collected and compared between groups.

### Effects of chronic mild unpredictable stress on instrumental learning and sensitivity to outcome devaluation

Male and female (Control: Final N = 22, 13 male; Stress: N = 25, 12 male) naïve mice were used in this experiment to assess how a recent history of chronic stress impacts instrumental learning and behavioral control strategy. 6 subjects (not included in above N) were excluded because they did not meet instrumental training performance criteria. Mice were randomly assigned to Control v. Stress groups. Mice were given 14 consecutive days of twice/daily stress or daily handling as described above. 24 hours after the final stress exposure, mice began instrumental conditioning as described above. After completion of FR-1, mice received 1 session each of training on an RR-2, RR-5, and RR-10 reinforcement schedule (max 20 outcomes/20 min/session). We chose an RR reinforcement schedule for this experiment because it tends to promote action-outcome learning and goal-directed decision making^96,97,102,104^ and would, thus, make it more difficult for prior stress to induce habits, increasing the robustness of the results. Following training, mice received a counterbalanced set of sensory-specific satiety outcome-specific devaluation tests, as above.

### Effects of chronic mild unpredictable stress on action-outcome learning

Male and female (Control, Non-degraded: Final N = 7, 3 male; Control, Contingency degradation N = 3, 3 male; Stress, Non-degraded: N = 7, 3 male; Stress, Contingency degradation N = 8, 4 male) naïve mice were used in this experiment assess how a recent history of chronic stress impacts the ability to learn an action-outcome contingency. 3 subjects (not included in above N) were excluded because they did not meet instrumental training performance criteria. Mice were randomly assigned to Control v. Stress groups. Mice were given 14 consecutive days of twice/daily stress or daily handling as described above. 24 hours after the final stress exposure, mice began instrumental conditioning as described above. After completion of FR-1, mice received 2 sessions of training in which lever presses were reinforced with a probability of 0.2 and one session in which they were reinforced with a probability of 0.1 (max 20 outcomes/20 min/session). Following training mice received a single contingency degradation or non-degraded control session, as described above. This was followed the next day by a lever-pressing probe test, described above.

### Effects of chronic mild unpredictable stress on common indices of anxiety- and depression-like behavior

Male and female (Control: Final N = 12, 6 male; Stress: N = 12, 6 male) naïve mice were used in this experiment to assess how a recent history of chronic stress impacts performance in common indices of anxiety- and depression-like behavior. Mice were randomly assigned to Control v. Stress groups. Mice were given 14 consecutive days of twice/daily stress or daily handling as described above. 24 hours after the final stress exposure, mice began testing, as described above. Mice were given tests in the order: open field test, light-dark emergence test, elevated plus maze, sucrose preference test, progressive ratio test.

### Tracing

Anterograde tracing of CeA neurons was performed as previously described^161^. Male (N = 2) and female (N = 2) naïve mice were infused bilaterally with the anterograde tracer AAV8-Syn-mCherry (Addgene, Watertown, MA) in the CeA (0.2 µL). Virus was allowed to express for 4 weeks, following which mice were perfused and histology was processed as described below to identify fluorescently labeled fibers in the dorsal striatum.

For retrograde tracing of DMS-projecting amygdala neurons, male (N = 2) and female (*N* = 2) naïve mice were infused with Fluorogold (Sigma, St. Louis, MO; 4% in sterile saline) in the DMS (0.2 µL). Virus was allowed to express for 5 days, following which mice were perfused and histology was processed as described below to identify fluorescently labeled cell bodies in CeA and BLA.

For retrograde tracing of monosynaptic inputs onto Drd1a^+^ or A2A^+^ DMS neurons, male (N = 4) and female (N = 4) *Drd1a-cre* or male (N = 3) and female (N = 2) *Adora2A-cre* naïve mice were infused with 0.3 µL AAV2-hSyn-FLEX-TVA-P2A-eGFP-2A-oG (Salk Gene Transfer, Targeting and Therapeutics Facility) in the DMS. Three weeks later, mice were infused with 0.3 µL EnvA G-deleted Rabies-mCherry at the same DMS coordinates. Mice were perfused 1 week later and tissue was processed as described below to identify monosynaptically-labeled inputs in CeA and BLA. 4 *Drd1a-cre* and 1 *Adora2A-cre* subjects were removed due to starter virus spillover in the BNST.

### Fiber photometry calcium imaging of CeA**→**DMS or BLA**→**DMS projections during instrumental learning following stress

Male and female (BLA→DMS Control: Final N = 9, 4 male; BLA→DMS Stress: N = 12, 5 male; CeA→DMS Control: N = 11, 6 male; CeA→DMS Stress: N = 11, 4 male) naïve mice were used in this experiment to monitor calcium fluctuations in CeA→DMS and BLA→DMS projections during instrumental conditioning after stress. 18 subjects (not included in above N) with off-target viral expression and/or fiber location were excluded from the dataset. 4 subjects were excluded for loss of optic fibers/headcaps. 4 subjects were excluded for missing recording data from one session. 3 subjects that did not complete instrumental conditioning were also excluded. Mice were randomly assigned to Virus and Stress groups. At surgery, mice received unilateral infusion (left/right hemisphere counterbalanced across subjects within each group) of a retrogradely trafficked AAV encoding cre-recombinase (AAVrg-Syn-Cre-P2A-dTomato, Addgene) into the DMS (0.3 µl) and of an AAV encoding the cre-dependent genetically encoded calcium indicator GCaMP8s (AAV9-Syn-FLEX-GcAMP8s-GFP, Addgene) into either the CeA or BLA (0.1-0.2 µl). Optic fiber cannulae (5.0-mm length (BLA) or 4.6 mm (CeA), 200-µm diameter, 0.37 NA, Inper, Hangzhou, China) were implanted over the GCaMP infusion site for calcium imaging at cell bodies. Mice were given 1 - 2 weeks to recover post-surgery, followed by 14 consecutive days of twice/daily stress or daily handling as described above. Mice were habituated to restraint during the final 3 days of the stress/handling period. 24 hours after the final stress exposure, mice began instrumental conditioning as described above. Each session began with a 3-minute baseline period prior to the start of the instrumental session for assessment of changes in baseline calcium activity. After completion of FR-1, mice received 1 session each of training on an RR-2, RR-5, and RR-10 reinforcement schedule (max 20 outcomes/20 min/session).

Fiber photometry was used to image bulk calcium activity in CeA→DMS or BLA→DMS neurons for 3-min prior to and throughout each instrumental conditioning session using a commercial fiber photometry system (Neurophotometrics Ltd., San Diego, CA). Two light-emitting LEDs (470 nm: Ca^2+^-dependent GCaMP fluorescence; 415 nm: autofluorescence, motion artifact, Ca^2+^-independent GCaMP fluorescence) were reflected off dichroic mirrors and coupled via a patch cord (200 µm; 0.37 NA, Inper) to the implanted optical fiber. The intensity of excitation light was adjusted to ∼100 µW at the tip of the patch cord. Fluorescence emission was passed through a 535-nm bandpass filter and focused onto the complementary metal-oxide semiconductor (CMOS) camera sensor through a tube lens. Samples were collected at 20 Hz interleaved between the 415 nm and 470 nm excitation channels using a custom Bonsai workflow. Time stamps of task events were collected simultaneously through an additional synchronized camera aimed at the Med Associates interface, which sent light pulses coincident with task events (onset, press, entry, reward). Signals were saved using Bonsai software and exported to MATLAB (MathWorks, Natick, MA) for analysis.

To assess the response to appetitive and aversive stimuli and provide a positive signal control, fiber photometry measurements were made during subsequent non-contingent reward and footshock sessions. In the first session, mice received 10 non-contingent food-pellet deliveries with a variable 60-s intertrial interval. 24 hours later, they received a session of 5, 2-s, 0.7mA footshocks with a variable 60-s intertrial interval. Calcium signal was aligned to reward collection or shock onset using timestamps collected as above. Mice were then perfused and brain tissue was processed with standard histology procedures described below to assess viral expression location/spread and fiber location.

#### Fiber photometry analysis

Data were pre-processed using a custom-written pipeline in MATLAB (MathWorks, Natick, MA) as previously^162^. The 415 nm and 470 nm signals were fit using an exponential curve. Change in fluorescence (ΔF/F) at each time point was calculated by subtracting the fitted 415 nm signal from the 470 nm signal and normalizing to the fitted 415 nm data [(470-fitted 415)/fitted 415)]. The ΔF/F data was Z-scored to the average of the whole session [(ΔF/F - mean ΔF/F)/std(ΔF/F)]. Z-scored traces were then aligned to behavioral event timestamps throughout each session. Area under the curve (AUC) was calculated for each individual aligned trace within each session using a trapezoidal function. We use the 3-s period prior to initiating presses to quantify activity related to the initiation of actions. We used the 3-s period following reward collection to quantify activity related to the earned outcome and unpredicted reward. We used the 1-s period following shock onset to quantify acute shock responses and the 2-s post-shock period to quantify activity following the shock. Quantifications and signal aligned to events were averaged across trials within a session and compared across sessions and between groups. Spontaneous activity was recorded during a 3-minute baseline period in the instrumental training context prior to each training session. Calcium events were identified as described previously^163^. First, we fitted the isosbestic channel to the 470 nm signal using an exponential function then subtracted the isosbestic trace from the calcium trace to remove calcium-independent artifacts. We defined a series of sliding-moving windows (15- s window, 1-second step) along the trace in which we filtered out high-amplitude events (greater than 2x the median of the 15-s window) and calculated the median absolute deviation of the resultant trace. Calcium transients with local maxima greater than 2 times above the median absolute deviation were selected as events. These events were used to calculate spontaneous event frequency and amplitude for BLA→DMS and CeA→DMS pathways.

### Optogenetic inhibition of BLA**→**DMS projections during instrumental learning

Male and female (eYFP Final N = 10, 5 male; Arch N = 11, 5 male) naïve mice were used in this experiment to assess the necessity of BLA→DMS projection activity at outcome experience during training for the action-outcome learning that supports goal-directed decision making. 13 subjects with off-target viral expression or fiber location and 6 subjects that did not complete instrumental conditioning were excluded from the dataset. Mice were randomly assigned to Viral group. At surgery, mice received bilateral infusion of an AAV encoding the inhibitory opsin archaerhodopsin (AAVDJ-Syn-eArch-YFP, Stanford Vector Core) or fluorophore control (AAVDJ-Syn-eYFP; Addgene) into the BLA (0.1 - 0.2 µl). Optic fiber cannulae (2.5-mm length, 100-µm diameter, 0.22 NA, Inper) were implanted over the DMS. Mice were given 3 weeks to recover and allow for viral expression. Mice were habituated to restraint for attaching optical fibers for 3 days immediately prior to instrumental conditioning. During instrumental conditioning, mice were tethered to a 100-µm diameter optic fiber bifurcated patchcord (Inper) attached to a 593-nm laser (Dragon Laser) via a rotary joint. Mice were habituated to the tether during the magazine training session, but no laser was delivered. Beginning with the first FR-1 session, all subjects received laser delivery during reward collection (first magazine entry after reward delivery; 5-s pulse, 8- 10 mW). After completion of FR-1, mice received 1 session each of instrumental conditioning on an RR-2, RR- 5, and RR-10 reinforcement schedule (max 20 outcomes/20 min/session). We chose an RR schedule of reinforcement for this experiment because tends to promote action-outcome learning and goal-directed decision making ^96,97,102,104^ and, thus, would make it more difficult to neurobiologically induce habit formation, increasing the robustness of the results. Following training mice received a counterbalanced set of sensory-specific satiety outcome-specific devaluation tests, as above. Mice were tethered but no laser was delivered on test days. Mice received laser as in training during the intervening retraining session. After instrumental training and testing, mice were tested in the RTPP test as described above. Mice were then perfused and brain tissue was processed with standard histology procedures described below to assess viral expression location/spread and fiber placement.

### Optogenetic activation of BLA**→**DMS projections during instrumental learning following stress

Male and female (Control eYFP: Final N = 11, 7 male; Control ChR2: N = 7, 4 males; Stress eYFP: N = 9, 2 male; Stress ChR2: N = 10 Stress, 3 male) mice were used in this experiment to assess whether activation of BLA→DMS projections during learning is sufficient to rescue action-outcome learning for goal-directed decision making in subjects with a history of stress. 4 subjects with off-target viral expression or fiber location and 2 subjects that did not complete instrumental conditioning were excluded from the dataset. Mice were randomly assigned to Virus and Stress groups. At surgery, mice received bilateral infusion of a retrogradely trafficked AAV encoding cre-recombinase (AAVrg-Syn-Cre-P2A-dTomato, Addgene) into the DMS (0.3 µl) and AAV encoding the cre-inducible excitatory opsin ChR2 (AAV8-Syn-DIO-ChR2-eYFP, Stanford Vector Core) or fluorophore control (AAV8-Syn-DIO-eYFP, Stanford Vector Core) into the BLA (0.1-0.2 µl). Optic fiber cannulae (5.0-mm length, 100-µm diameter, 0.22 NA, Inper) were implanted over the BLA. Mice were given 1 - 2 weeks to recover post-surgery, followed by 14 consecutive days of twice/daily stress or daily handling as described above. Mice were habituated to restraint for attaching optical fibers during the final 3 days of the stress/handling period. 24 hours after the final stress exposure, mice began instrumental conditioning, as described above. During instrumental conditioning, mice were tethered to a 100-µm diameter optic fiber bifurcated patchcord (Inper) attached to a 473-nm laser (Dragon Laser) via a rotary joint. Mice were habituated to the tether during the magazine training session, but no laser was delivered. Beginning with the first FR-1 session, all subjects received laser delivery during reward collection, (first magazine entry after reward delivery; 2-s duration, 20 Hz, 5-ms pulse width, 8-10 mW). After completion of FR-1, mice received 1 training session on an RI-15s reinforcement schedule and 2 training sessions on the RI-30s schedule (max 20 outcomes/20 min/session). We chose an RI reinforcement schedule for this experiment because it tends to promote habit formation ^96,97,102,104^ and, thus, would make it more difficult to neurobiologically prevent stress-potentiated habit, increasing the robustness of the results. Following training, mice received a counterbalanced set of sensory-specific satiety outcome-specific devaluation tests, as above. Mice were tethered but no laser was delivered on test days. Mice received laser as in training during the intervening retraining session. Mice were then perfused and brain tissue was processed with standard histology procedures described below to assess viral expression location/spread and fiber placement.

### Chemogenetic activation of BLA**→**DMS projections during instrumental learning following stress

Male and female (Control, mCherry: Final N = 12, 7 male; Control, hM3Dq: N = 6, 3 male; Stress, mCherry: N = 9, 5 male; Stress, hM3Dq: N = 10, 5 male) naïve mice were used in this experiment to assess whether activation of BLA→DMS projections during learning is sufficient to rescue action-outcome learning for goal-directed decision making in subjects with a history of stress. 15 subjects with off-target viral expression and 2 subjects that did not complete instrumental conditioning were excluded from the dataset. Mice were randomly assigned to Viral and Stress groups. At surgery, all mice received bilateral infusion of the retrogradely-trafficked canine- adenovirus encoding cre-recombinase (CAV2-Cre-GFP; Plateforme de Vectorologie de Montpellier, Montpellier, France) into the DMS (0.3 µl) and AAV encoding the cre-inducible excitatory designer receptor human M3 muscarinic receptor (hM3Dq; AAV2-Syn-DIO-hM3Dq-mCherry; Addgene, Watertown, MA) or fluorophore control (AAV2-Syn-DIO-mCherry; Addgene) into the BLA (0.1-0.2 µl). Mice were given 1-2 weeks to recover post-surgery, followed by 14 consecutive days of twice/daily stress or daily handling, as described above. Mice were habituated to i.p. injections during the final 3 days of the stress/handling period. 24 hours after the final stress exposure, mice began instrumental conditioning, as described above. All subjects received an intraperitoneal (i.p.) injection of clozapine-*n*-oxide (water soluble CNO; 0.2 mg/kg^105–108,164^; Hello Bio, Princeton, NJ) 30 min prior to each instrumental conditioning session. Upon completion of FR-1 (80% max rewards delivered), mice received 1 training session the RI-15s reinforcement schedule following by 2 sessions on an RI-30s schedule (max 30 outcomes/30 min/session). We chose an RI schedule of reinforcement for this experiment because it tends to promote habit formation ^96,97,102,104^ and, thus, would make it more difficult to neurobiologically prevent stress-potentiated habit, increasing the robustness of the results. Following training mice received a counterbalanced pair of sensory-specific satiety outcome-specific devaluation tests, as above. No CNO was given on test days. CNO was given prior to the retraining session (RI-30s) in between tests. After instrumental training and testing, mice were perfused and brain tissue was processed using standard histology procedures described below to assess viral expression location and spread.

### Optogenetic inactivation of CeA**→**DMS projections during instrumental overtraining

Male and female (control, eYFP: N = 11, 3 male; control, Arch: N = 11, 7 male) naïve mice were used in this experiment to assess the necessity of CeA→DMS projection activity at outcome experience during learning for the natural habit formation that occurs with overtraining. 2 subjects with off-target viral expression or fiber location were excluded from the dataset. Mice were randomly assigned to Virus group. At surgery, mice received bilateral infusion of an AAV encoding the inhibitory opsin Arch (AAVDJ-Syn-eArch-eYFP, Stanford Vector Core) or fluorophore control (AAVDJ-Syn-eYFP; Addgene) into the CeA (0.1-0.2 µl). Optic fiber cannulae (2.5-mm length, 100-µm diameter, 0.22 NA, Inper) were implanted over the DMS. Mice were given 1 week to recover post-surgery. Mice were habituated to restraint for attaching optical fibers. Mice then receive instrumental overtraining on the RI-30s schedule as described above. During instrumental conditioning, mice were tethered to a 100-µm diameter optic fiber bifurcated patchcord (Inper) attached to a 593-nm laser (Dragon Laser) via a rotary joint. Mice were habituated to the tether during the magazine training session, but no laser was delivered. Beginning with the first FR-1 session, all subjects received laser delivery during reward collection, (first magazine entry after reward delivery; 5-s pulse, 8-10 mW). After completion of FR-1, mice received 1 training session on an RI- 15s reinforcement schedule and 7 training sessions on the RI-30s schedule (max 20 outcomes/20 min/session). We chose an RI reinforcement schedule for this experiment because it tends to promote habit formation ^96,97,102,104^. We overtrained subjects to also promote the formation of habits naturally in control subjects. Following training, mice received a counterbalanced set of sensory-specific satiety outcome-specific devaluation tests, as above. Mice were tethered but no laser was delivered on test days. Mice received laser as in training during the intervening retraining session. Mice were then perfused and brain tissue was processed with standard histology procedures described below to assess viral expression location/spread and fiber placement.

### Optogenetic inactivation of CeA**→**DMS projections during instrumental learning following stress

Male and female (Control, eYFP: N = 9, 5 male; Control, Arch: N = 11, 4 male; Stress, eYFP: N = 7, 6 male; Stress, Arch: N = 9, 5 male) naïve mice were used in this experiment to assess the necessity of CeA→DMS projection activity at outcome experience during learning for stress-potentiated habit formation. 12 subjects with off-target viral expression or fiber location and 2 subjects that did not complete instrumental conditioning were excluded from the dataset. Mice were randomly assigned to Virus and Stress groups. At surgery, mice received bilateral infusion an AAV encoding the inhibitory opsin Arch (AAVDJ-Syn-eArch-eYFP, Stanford Vector Core) or fluorophore control (AAVDJ-Syn-eYFP; Addgene) into the CeA (0.1-0.2 µl). Optic fiber cannulae (2.5-mm length, 100-µm diameter, 0.22 NA, Inper) were implanted over the DMS. Mice were given 1 - 2 weeks to recover post-surgery, followed by 14 consecutive days of twice/daily stress or daily handling as described above. Mice were habituated to restraint for attaching optical fibers during the final 3 days of the stress/handling period. 24 hours after the final stress exposure, mice began instrumental conditioning as described above. During instrumental conditioning, mice were tethered to a 100-µm diameter optic fiber bifurcated patchcord (Inper) attached to a 593- nm laser (Dragon Laser) via a rotary joint. Mice were habituated to the tether during the magazine training session, but no laser was delivered. Beginning with the first FR-1 session, all subjects received laser delivery during reward collection, (first magazine entry after reward delivery; 5-s pulse, 8-10 mW). After completion of FR-1, mice received 1 training session on an RI-15s reinforcement schedule and 2 training sessions on the RI- 30s schedule (max 20 outcomes/20 min/session). We chose an RI reinforcement schedule for this experiment because it tends to promote habit formation ^96,97,102,104^ and, thus, would make it more difficult to neurobiologically prevent stress-potentiated habit, increasing the robustness of the results. Following training, mice received a counterbalanced set of sensory-specific satiety outcome-specific devaluation tests, as above. Mice were tethered but no laser was delivered on test days. Mice received laser as in training during the intervening retraining session. After instrumental training and testing, mice were tested in the RTPP test as described above. Mice were then perfused and brain tissue was processed with standard histology procedures described below to assess viral expression location/spread and fiber placement.

### Chemogenetic inactivation of CeA**→**DMS projections during instrumental learning following stress

Male and female (Control, mCherry: N = 12, 5 male; Control, hM4Di: N = 13, 8 male; Control, mCherry: N = 11, 5 male; Control, hM4Di: N = 9, 4 male) naïve mice were used in this experiment to assess the necessity of CeA→DMS projection activity during learning for stress-potentiated habit formation. 16 subjects with off-target viral expression and 3 subjects that did not complete instrumental conditioning were excluded from the dataset. Mice were randomly assigned to Viral and Stress groups. At surgery, all mice received bilateral infusion of the retrogradely-trafficked CAV encoding cre-recombinase (CAV2-Cre-GFP; Plateforme de Vectorologie de Montpellier) into the DMS (0.3 µl) and AAV encoding the cre-inducible inhibitory designer receptor human M4 muscarinic receptor (hM4DGi; AAV2-Syn-DIO-hM4Di-mCherry; Addgene) or fluorophore control (AAV2-Syn-DIO-mCherry; Addgene) into the CeA (0.1-0.2 µl). Mice were given 1-2 weeks to recover post-surgery, followed by 14 consecutive days of twice/daily stress or daily handling as described above. Mice were habituated to i.p. injections during the final 3 days of the stress/handling period. 24 hours after the final stress exposure, mice began instrumental conditioning as described above. All subjects received an intraperitoneal (i.p.) injection of CNO (2 mg/kg^111,112,164,165^; Hello Bio) 30 min prior to each instrumental conditioning session. Upon completion of FR-1, mice received 1 session of training on an RI-15s reinforcement schedule followed by 2 sessions on the RI-30s schedule (max 30 outcomes/30 min/session). We chose an RI reinforcement schedule for this experiment because it tends to promote habit formation ^96,97,102,104^ and, thus, would make it more difficult to neurobiologically prevent stress-potentiated habit, increasing the robustness of the results. Following training, mice received a counterbalanced pair of sensory-specific satiety outcome-specific devaluation tests, as above. No CNO was given on test days. CNO was given prior to the retraining session. After instrumental training and testing, mice were then perfused and brain tissue was processed with standard histology procedures described below to assess viral expression location and spread.

### Optogenetic activation of CeA**→**DMS projections during instrumental learning

Male and female (eYFP *N* = 17, 9 male; ChR2: *N* = 6, 3 male) naïve mice were used in this experiment to assess whether CeA→DMS projection activation at outcome experience during learning is sufficient to promote habit formation. 11 subjects with off-target viral expression or fiber location and 4 subjects that did not complete instrumental conditioning were excluded from the dataset. Mice were randomly assigned to Virus group. Given the low density of CeA→DMS projections, we choose to activate DMS-projecting CeA cell bodies. At surgery, mice received bilateral infusion of a retrogradely trafficked AAV encoding cre-recombinase (AAVrg-Syn-Cre- P2A-dTomato, Addgene) into the DMS (0.3 µl) and AAV encoding the cre-inducible excitatory opsin channelrhodopsin 2 (ChR2; AAV8-Syn-DIO-ChR2-eYFP, Stanford Vector Core) or fluorophore control (AAV8- Syn-DIO-eYFP, Stanford Vector Core) into the CeA (0.1-0.2 µl). Optic fiber cannulae (5.0-mm length, 100-µm diameter, 0.22 NA, Inper) were implanted over the CeA. Mice were given 3 weeks to recover and allow for viral expression. Mice were habituated to restraint for 3 days prior to instrumental conditioning. During instrumental conditioning, mice were tethered to a 100-µm diameter optic fiber bifurcated patchcord (Inper) attached to a 473- nm laser (Dragon Laser) via a rotary joint. Mice were habituated to the tether during the magazine training session, but no laser was delivered. Beginning with the first FR-1 session, all subjects received laser delivery during reward collection (first magazine entry after reward delivery; 2-s duration, 20 Hz, 5-ms pulse width, 8-10 mW). After completion of FR-1, mice received 1 day each of training on an RR-2, RR-5, and RR-10 reinforcement schedule (max 20 outcomes/20 min/session). We chose an RR schedule of reinforcement for this experiment because it tends to promote action-outcome learning and goal-directed decision making ^96,97,102,104^ and, thus, would make it more difficult to neurobiologically induce habit formation, increasing the robustness of the results. Following training mice received a counterbalanced set of sensory-specific satiety outcome-specific devaluation tests, as above. Mice were tethered but no laser was delivered on test days. Mice received laser as in training during the intervening retraining session. After instrumental training and testing, mice were tested in the RTPP test, as described above. Mice were then perfused and brain tissue was processed with standard histology procedures described below to assess viral expression location/spread and fiber placement.

### Optogenetic activation of CeA**→**DMS projections during instrumental learning following sub-threshold stress

Male and female (eYFP N = 10, 4 male; ChR2: N = 12, 6 male) naïve mice were used in this experiment to assess whether CeA→DMS projection activation at outcome experience during learning is sufficient to promote habit formation in mice with a history of less-frequent stress (subthreshold for promoting habit formation). 11 subjects with off-target viral expression or fiber location and 4 subjects that did not complete instrumental conditioning were excluded from the dataset. Mice were randomly assigned to viral groups. Similar to optogenetic activation of CeA→DMS neurons in control mice, we chose to target cell bodies with this approach. At surgery, mice received bilateral infusion of a retrogradely trafficked AAV encoding cre-recombinase (AAVrg-Syn-Cre- P2A-dTomato, Addgene) into the DMS (0.3 µl) and AAV encoding the cre-inducible excitatory opsin ChR2 (AAV8-Syn-DIO-ChR2-eYFP, Stanford Vector Core) or fluorophore control (AAV8-Syn-DIO-eYFP, Stanford Vector Core) into the CeA (0.1-0.2 µl). Optic fiber cannulae (5.0-mm length, 100-µm diameter, 0.22 NA, Inper) were implanted over the CeA. Mice were given 1 - 2 weeks to recover post-surgery, followed by 14 consecutive days of once/daily stress or daily handling as described above. Mice were habituated to restraint for attaching optical fibers during the final 3 days of the subthreshold stress/handling period. 24 hours after the final stress exposure, mice began instrumental conditioning, as described above. During instrumental conditioning, mice were tethered to a 100-µm diameter optic fiber bifurcated patchcord (Inper) attached to a 473-nm laser (Dragon Laser) via a rotary joint. Mice were habituated to the tether during the magazine training session, but no laser was delivered. Beginning with the first FR-1 session, all subjects received laser delivery during reward collection, (first magazine entry after reward delivery; 2-s duration, 20 Hz, 5-ms pulse width, 8-10 mW). After completion of FR-1, mice received 1 day each of training on an RR-2, RR-5, and RR-10 reinforcement schedule (max 20 outcomes/20 min/session). Following training mice received a counterbalanced set of sensory-specific satiety outcome-specific devaluation tests, as above. Mice were tethered but no laser was delivered on test days. Mice received laser during the intervening retraining session. After instrumental training and testing, mice were tested in the RTPP test, as described above. Mice were then perfused and brain tissue was processed with standard histology procedures described below to assess viral expression location/spread and fiber placement.

### Immunohistochemistry

Mice were anesthetized with isoflurane and transcardially perfused with ice-cold PBS followed by cold 4% paraformaldehyde. The brains were removed, post-fixed in 4% paraformaldehyde, then cryoprotected in 30% sucrose in PBS. 30-µm coronal slices were taken on a cryostat and collected in PBS. Immunohistochemical analysis was performed as described previously^162,166–168^. Briefly, floating sections were blocked for 1 hour at room temperature in blocking solution (3% normal goat serum (NGS, Jackson ImmunoResearch Laboratories), 0.3% Triton X-100 (Fisher)) in PBS and then incubated overnight with gentle agitation at 4 °C in blocking solution plus 1:1000 dilution primary antibody (chicken anti-GFP polyclonal, Abcam; rabbit anti-dsRed polyclonal, Takara Bio). Sections were then incubated covered with gentle agitation for 2 hours at room temperature in blocking solution plus 1:500 dilution secondary antibody (goat anti-rabbit IgG Alexafluor 594 conjugate; goat anti-chicken IgG Alexafluor 488 conjugate, Invitrogen). All sections were washed 3 times for 5 min each in PBS before and after each incubation step and mounted on slides using ProLong Gold antifade reagent with DAPI (Invitrogen). All images were acquired using a Keyence (BZ-X710) microscope with 4X, 10X, and 20X objectives (CFI Plan Apo), CCD camera, and BZ-X Analyze software and a Zeiss Confocal LSM with 2.5X and 20X objectives and Zeiss ZEN (blue edition) image acquisition software.

### Statistical Analysis

Datasets were analyzed by 2-tailed t-tests, or 1-, 2-, or 3-way repeated-measures analysis of variance (ANOVA), as appropriate (GraphPad Prism, GraphPad, San Diego, CA; SPSS, IBM, Chicago, IL). For chemogenetic replications of optogenetic results, we used planned comparisons for test press rate data. Some datasets were slightly non-normally distributed. For these datasets, statistical tests were also run using non-parametric analyses and the results were highly consistent. We opted to use parametric statistics for consistency across experiments and given evidence that ANOVA is robust to slight non-normality^169,170^. Bonferroni post hoc tests corrected for multiple comparisons were performed to clarify statistical interactions. Greenhouse-Geisser correction was applied to mitigate the influence of unequal variance between conditions. Alpha levels were set at *P* < 0.05.

### Sex as a biological variable

For the initial behavioral finding, sex was included as a factor in the ANOVA and found to not significantly account for variance (No main effect of Sex on lever pressing acquisition: F_(1, 43)_ = 0.43, *P* = 0.51, devaluation test press rate: F_(1,43)_ = 0.60, *P* = 0.44, or devaluation index: F_(1,43)_ = 0.04, *P* = 0.84). Therefore, data from male and female mice was combined for analyses. For subsequent experiments, male and female mice were used in approximately equal numbers, but the N per sex was underpowered to examine sex differences. Sex was therefore not included as a factor in statistical analyses, though individual data points are visually disaggregated by sex.

### Rigor and reproducibility

Group sizes were estimated based on prior work with this behavioral task^171^ and to ensure counterbalancing of virus, stress, pellet type, and devaluation test order. Investigators were not blinded to viral or stress group because they were required to administer infusions and stress exposure. All behaviors were scored using automated software (Med Associates). Each experiment included at least 1 replication cohort and cohorts were balanced by Viral group, Stress group, and hemisphere (for photometry recordings and tracing) prior to the start of the experiment. Investigators were blinded to group when performing histological validation and determining exclusions based on viral spread or mistargeted implant.

### Data and code availability

All data that support the findings of this study are available in the source data accompanying this paper and from the corresponding author upon request. Custom-written MATLAB code is accessible via Dryad repository^172^ and available from the corresponding author upon request.

## EXTENDED DATA FIGURES

**Extended data Figure 1-1:**
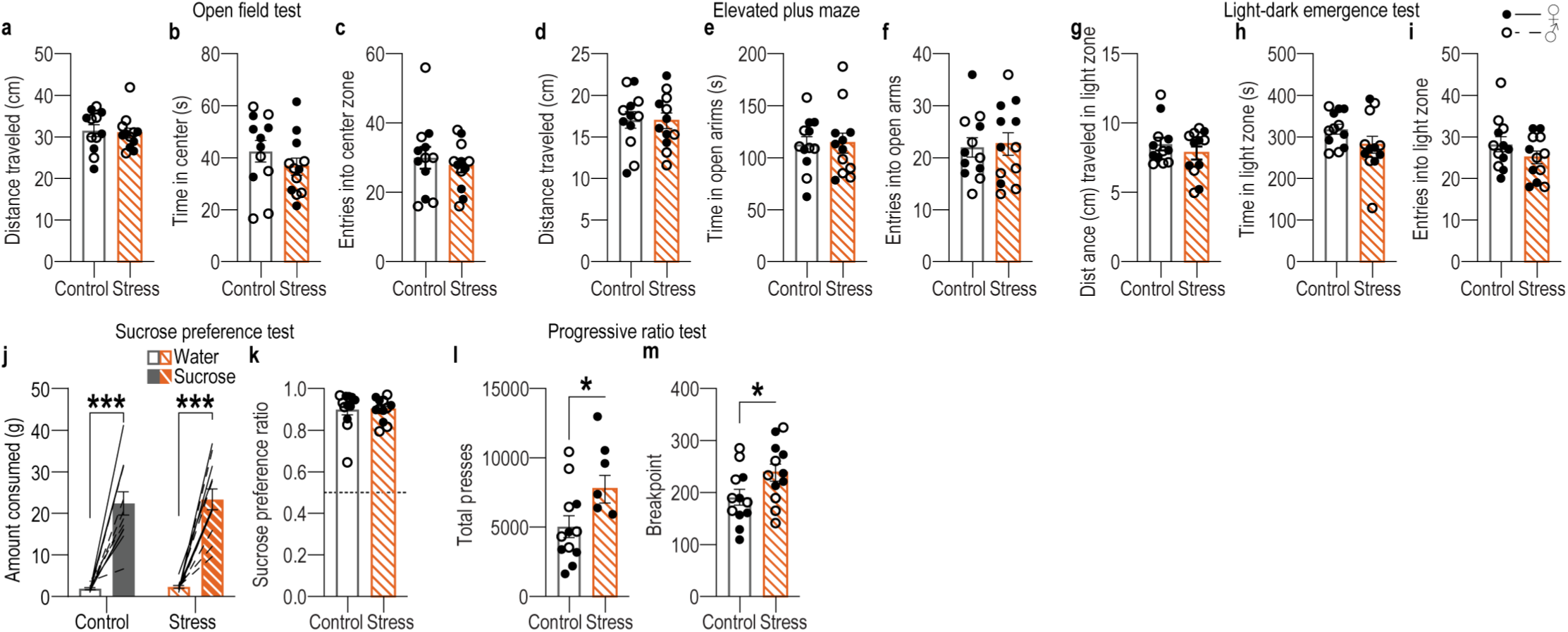
Chronic mild unpredictable stress does not cause classic anxiety- and depression-like phenotypes. Mice received 14 consecutive d of chronic mild unpredictable stress (stress) including twice daily exposure to 1 of 6 mild stressors at pseudorandom times and orders: damp bedding (16 hr), tilted cage (16 hr), white noise (80 db; 2 hr), continuous illumination (8 hr), physical restraint (2 hr), footshock (0.7- mA, 1-s, 5 shocks/10 min) prior to subsequent testing in a battery of behavioral assays classically used to assess anxiety- and depression-like behavior. **(a-c)** Open field test. Distance traveled (a; t_(22)_ = 0.32, *P* = 0.75), time spent in center zone (b; t_(22)_ = 1.10, *P* = 0.28), and entries into center zone (c; t_(22)_ = 0.63, *P* = 0.54). **(d-f)** Elevated plus maze. Distance traveled (d; t_(22)_ = 0.08, *P* = 0.94), time spent in open arms (e; t_(22)_ = 0.01, *P* = 0.92), and entries into open arms (f; t_(22)_ = 0.23, *P* = 0.82). **(g-i)** Light-dark emergence test. Distance traveled in light zone (g; t_(22)_ = 0.97, *P* = 0.34), time spent in light zone (h; t_(22)_ = 1.57, *P* = 0.13), and entries into light zone (I; t_(22)_ = 1.37, *P* = 0.19). **(j-k)** Sucrose preference test. Average amount consumed of water and 10% sucrose over 24 hr (j; Solution: F_(1, 22)_ = 113.20, *P* < 0.0001; Stress: F_(1, 22)_ = 0.14, *P* = 0.71, Solution x Stress: F_(1, 22)_ = 0.02, P = 0.89) and ratio of sucrose:water consumed (k; t_(22)_ = 0.03, *P* = 0.98). **(l-m)** Progressive ratio Tests. Total presses (l; t_(22)_ = 2.13, *P* = 0.04) and breakpoint (k; Final ratio completed; t_(22)_ = 2.12, *P* = 0.46). Control N = 12 (6 male), Stress N = 12 (6 male). Males = closed circles, Females = open circles. \**P* <0.05, \*\*\**P* <0.001. Our stress procedure does not affect general locomotor activity or avoidance of anxiogenic spaces or create an anhedonia phenotype. Rather this stress procedure appears to cause elevated motivation to exert effort to obtain reward. This contrasts with more severe, longer-lasting stress procedures, which do produce anxiety- and depression-like phenotypes in these tasks^155,173,174^. Thus, our stress procedure models chronic, low-level stress.

**Extended data Figure 1-2:**
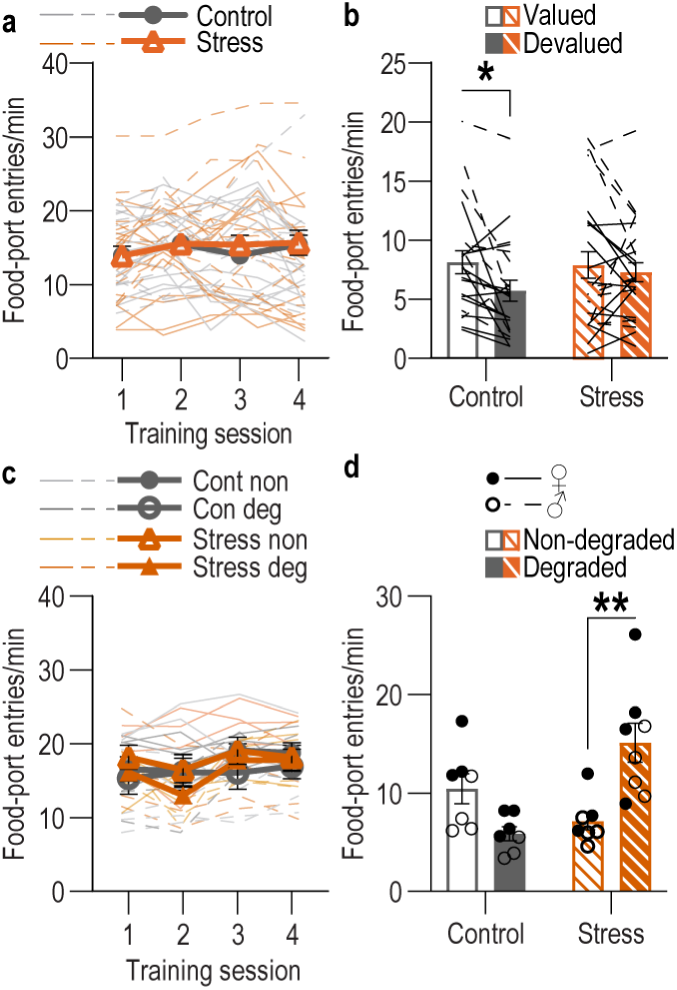
Food-port entries during training and probe tests following handling control or chronic stress. **(a)** Food-port entry rate across training for subjects in the devaluation experiment. Training: F_(2.42, 108.90)_ = 3.17, *P* = 0.04; Stress: F_(1, 45)_ = 0.07, *P* = 0.79; Training x Stress: F_(3, 135)_ = 0.57, *P* = 0.64. **(b)** Food-port entries during the devaluation probe tests. Value: F_(1, 45)_ = 6.77, *P* = 0.01, Stress: F_(1, 45)_ = 0.29, *P* = 0.60; Stress x Value: F_(1, 45)_ = 2.42, *P* = 0.13. Control N = 22 (13 male), Stress N = 25 (12 male). **(c)** Food-port entry rate across training for subjects in the contingency degradation experiment. Training: F_(2.84, 62.10)_ = 6.44, *P* = 0.001; Stress: F_(1, 25)_ = 0.01, *P* = 0.91; Future Contingency Degradation group: F_(1, 25)_ = 1.27, *P* = 0.27; Training x Stress: F_(3, 75)_ = 1.62, *P* = 0.19; Training x Group: F_(3, 75)_ = 0.24, *P* = 0.87; Stress x Group: F_(1, 25)_ = 0.004, *P* = 0.95; Training x Stress x Group: F_(3, 75)_ = 1.49, *P* = 0.23. **(d)** Food-port entries during the probe test 24 hr following contingency degradation or non-degraded control. Stress x Contingency Degradation Group: F_(1, 25)_ = 18.88, *P* = 0.0002; Contingency Degradation: F_(1, 25)_ = 4.29, *P* = 0.05; Stress: F_(1, 25)_ = 1.41, *P* = 0.25. Control, Non-degraded N = 7 (3 male), Control, Degraded N = 7 (3 male), Stress Non-degraded N = 7 (3 male) Stress Degraded N = 8 (4 male). Males = solid lines, Females = dashed lines. \**P* < 0.05, \*\**P* < 0.01.

**Extended data Figure 1-3:**
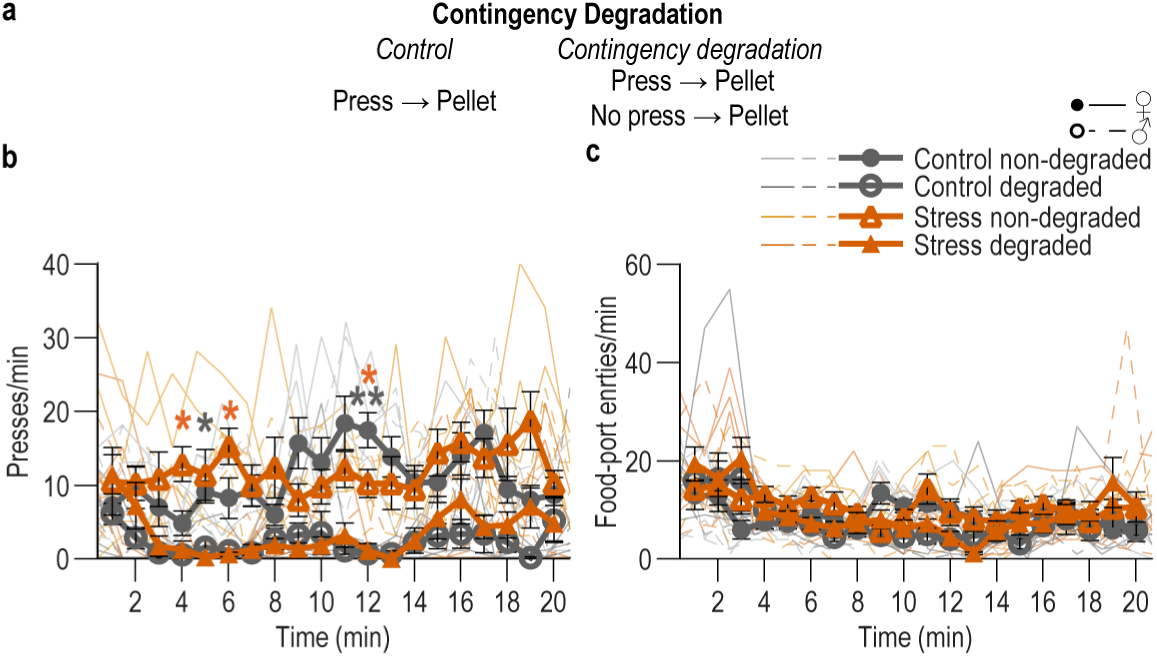
Lever presses and food-port entries during contingency degradation. **(a)** Contingency degradation procedure schematic. Following stress and training, half the subjects in each group received a 20-min contingency degradation session during which lever pressing continued to earn reward with a probability of 0.1, but reward was also delivered non-contingently with the same probability. This session was identical for non-degraded controls, except they did not receive free rewards. **(b)** Press rate in 1-min bins during the contingency degradation session. Time x Contingency Degradation Group: F_(19, 475)_ = 2.03, *P* = 0.0063; Time x Stress: F_(19, 475)_ = 2.43, *P* = 0.0007; Stress x Group: F_(1, 25)_ = 0.0001, *P* = 0.99; Time: F_(9.17, 229.20)_ = 2.13, *P* = 0.03; Stress: F_(1, 25)_ = 1.36, *P* = 0.26; Degradation Group: F_(1, 25)_ = 68.23, *P* < 0.0001; Time x Stress x Degradation Group: F_(19, 475)_ = 1.30, *P* = 0.19. Contingency degradation cause lower press rates across the session in both control (Time x Contingency Degradation Group: F_(12, 228)_ = 2.47, *P* = 0.0009; Time: F_(6.62, 79.39)_ = 2.47, *P* = 0.03; Degradation Group: F_(1, 12)_ = 45.16, *P* < 0.0001) and stressed (Contingency Degradation Group: F_(1, 13)_ = 28.22, *P* = 0.0001; Time: F_(6.01, 78.16)_ = 2.19, *P* = 0.05; Time x Contingency Degradation Group: F_(19, 247)_ = 1.10, *P* = 0.35) mice. **(c)** Rate of entry into the food-delivery port in 1-min bins during the contingency degradation session. Time x Contingency Degradation Group: F_(19, 475)_ = 3.80, *P* < 0.0001; Time x Stress: F_(19, 475)_ = 1.20, *P* = 0.26; Stress x Group: F_(1, 25)_ = 0.006, *P* = 0.94; Time: F_(6.26, 156.60)_ = 7.53, *P* < 0.0001; Stress: F_(1, 25)_ = 2.51, *P* = 0.13; Degradation Group: F_(1, 25)_ = 1.37, *P* = 0.5; Time x Stress x Degradation Group: F_(19, 475)_ = 0.86, *P* = 0.63. Control, Non-degraded N = 7 (3 male), Control, Degraded N = 7 (3 male), Stress Non-degraded N = 7 (3 male) Stress Degraded N = 8 (4 male). Males = closed circles/solid lines, Females = open circles/dashed lines. \**P* <0.05, \*\**P* < 0.01.

**Extended Data Figure 2-1:**
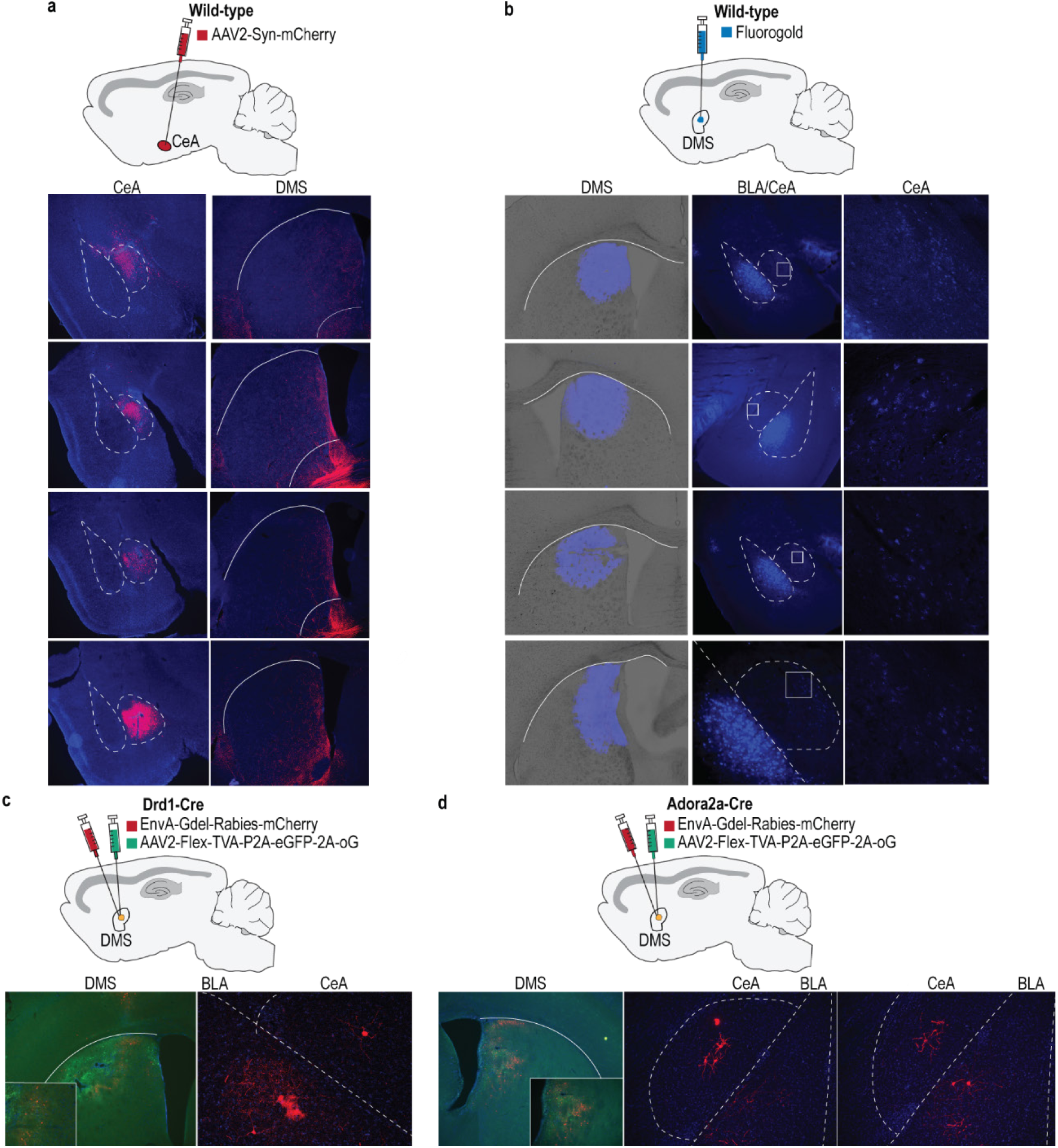
BLA and CeA directly project to DMS. **(a)** Top: Anterograde tracing approach. Infusion of an AAV expressing mCherry into the CeA. Bottom: mCherry labeling at infusion site in CeA (left) and mCherry-labeled fibers in the DMS (right). N = 4 (2 male). We observed mCherry-expressing putative fibers in the DMS but not dorsolateral striatum. Expression was also detected in other well-known CeA projection targets such as the bed nucleus of the stria terminalis. **(b)** Top: Retrograde tracing approach. We infused the fluorescently labeled retrograde tracer Fluorogold into the DMS. Bottom: Fluorogold labeling at infusion site in DMS (left) and fluorogold-labeled, DMS-projecting cell bodies in BLA and CeA (middle), with CeA magnified (right). Labeled cells was detected in both BLA and CeA, indicating that both BLA and CeA directly project to DMS. Labeling was greater in BLA than CeA, indicating the BLA→DMS pathway is denser than the CeA→DMS pathway. N = 4 (2 male). **(c)** Top: Approach for rabies trans-synaptic retrograde tracing of DMS Drd1^+^ striatal neurons. We used rabies tracing to confirm monosynaptic amygdala projections onto DMS neurons. We infused a starter virus expressing cre-dependent TVA-oG-GFP into the DMS of mice expressing cre-recombinase under the control of dopamine receptor 1 (D1-Cre) or adenosine 2a receptor (A2A-Cre) genes^175,176^, followed by ΔG- deleted rabies-mCherry to retrogradely label cells that synapse onto DMS D1 or A2A neurons. Bottom: Starter oG virus (green) and ΔG-deleted rabies-mCherry (red) expression in DMS Drd1^+^ neurons (left) and rabies-labeled, DMS D1-projecting cell bodies in the BLA and CeA (right), consistent with prior reports^80,81^. Representative example from N = 4 (3 males). **(d)** Top: Approach for rabies trans-synaptic retrograde tracing of DMS Adora2a^+^ neurons. Bottom: Starter ΔG virus (green) and rabies-mCherry (red) expression in DMS Adora2a^+^ neurons (left) and rabies-labeled, DMS A2A-projecting cell bodies in the BLA and CeA (right). Representative example N = 4 (3 males). Combined, these data confirm that both BLA and CeA directly project to the DMS and are, thus, poised to influence the learning that supports goal-directed decision making and habit formation.

**Extended Data Figure 2-2:**
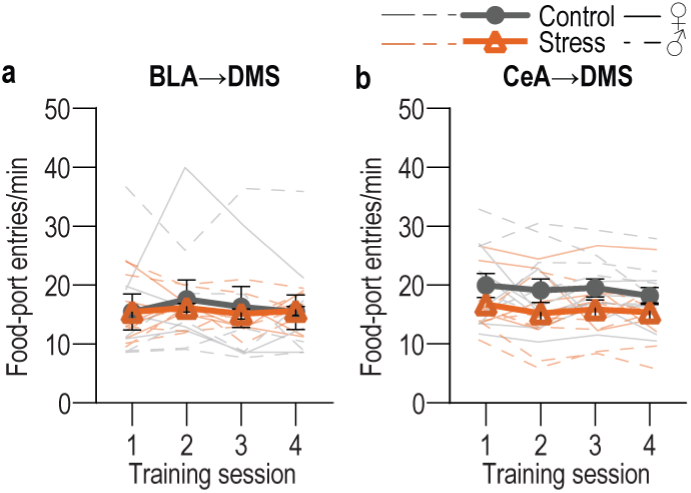
Food-port entries during training with fiber photometry recording of BLA→DMS or CeA→DMS calcium activity following handling control or chronic stress. **(a)** Food-port entry rates across training for BLA→DMS GCaMP8s mice. Training: F_(2.47, 46.99)_ = 0.65, *P* = 0.56; Stress: F_(1, 19)_ = 0.05, *P* = 0.82; Training x Stress: F_(3, 57)_ = 0.24, *P* = 0.87. BLA Control N = 9 (4 male), BLA Stress N = 12 (5 male). **(b)** Food-port entry rates across training for CeA→DMS GCaMP8s mice. Training: F_(2.36, 47.19)_ = 0.89, *P* = 0.43; Stress: F_(1, 20)_ = 2.71, *P* = 0.12; Training x Stress: F_(3, 60)_ = 0.09, *P* = 0.96. CeA Control N = 11 (6 male), CeA Stress N = 11 (4 male). Males = solid lines, Females = dashed lines.

**Extended Data Figure 2-3:**
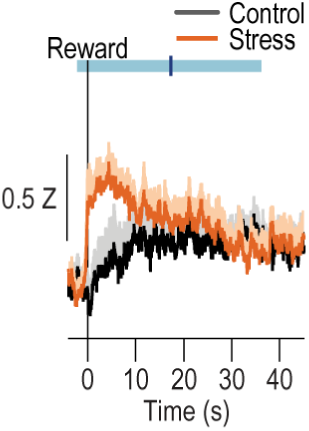
CeA→DMS pathway activity following reward collection. Trial-averaged Z-scored Δf/F CeA→DMS GCaMP8s fluorescence changes aligned to reward collection, with 40-s post-collection window. Shading reflects between-subject s.e.m. Blue line is the average time of the next lever press (light blue bar = s.e.m.). In stressed mice, CeA→DMS neurons respond to earned reward and this activity takes ∼30 s on average to come back to baseline. Control N = 11 (6 male), Stress N = 11 (4 male).

**Extended Data Figure 2-4:**
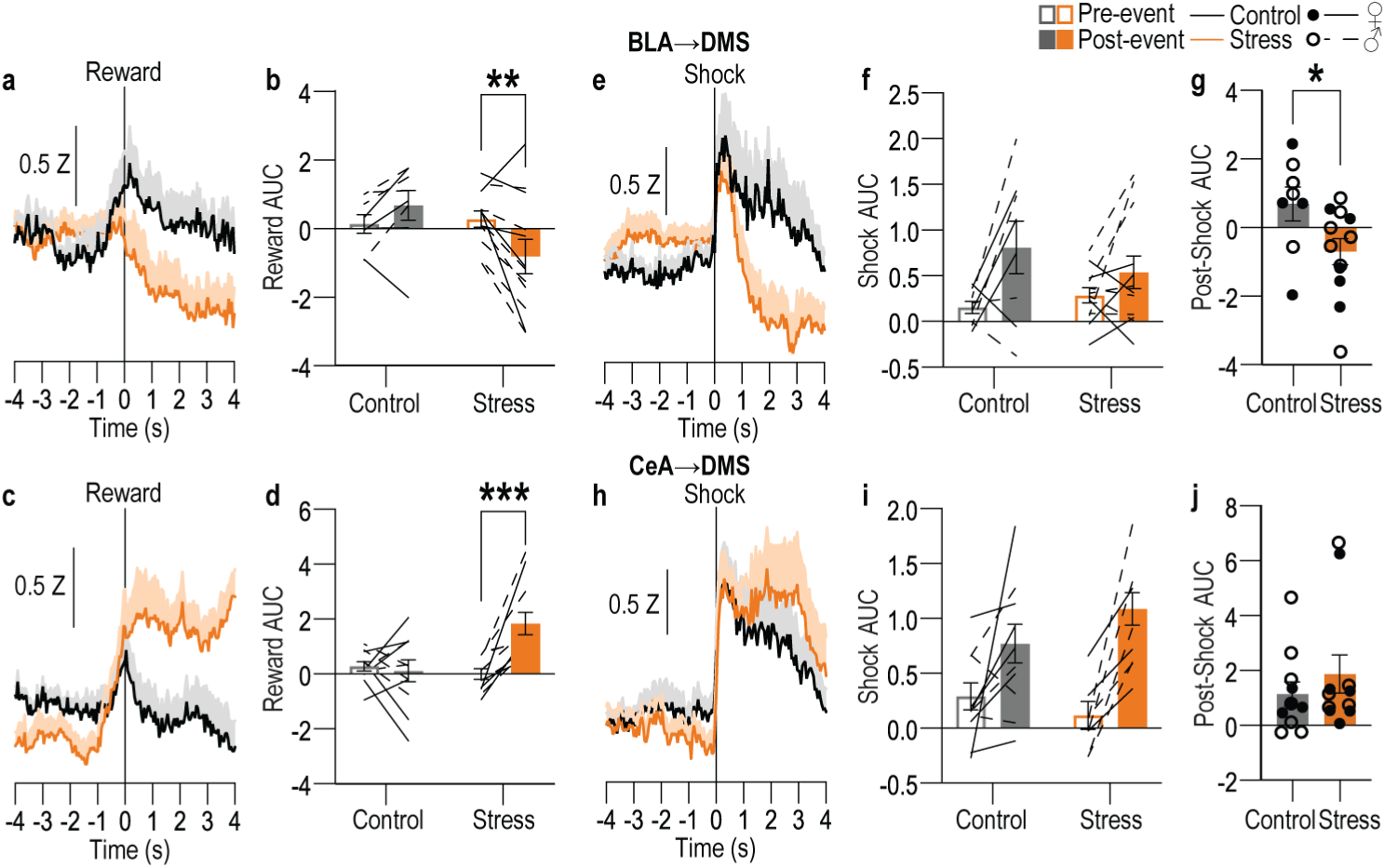
BLA→DMS and CeA→DMS pathway responses to unpredicted rewarding and aversive events in control and stressed mice. Following instrumental training (Figure 2), we used fiber photometry to record GCaMP8s fluorescent changes in either BLA (top) or CeA (bottom) neurons that project to the DMS in response to unpredicted food-pellet reward deliveries or unpredicted 2-s, 0.7 mA footshocks in control and stressed mice. **(a)** Trial-averaged Z-scored Δf/F BLA→DMS GCaMP8s fluorescence changes around unpredicted food-pellet reward delivery. **(b)** Trial-averaged quantification of area under the BLA→DMS GCaMP8s Z-scored Δf/F curve (AUC) during the 3-s period prior to (baseline) and following reward collection. Stress x Reward: F_(1, 18)_ = 10.88, *P* = 0.004; Reward: F_(1, 18)_ = 1.19; *P* = 0.03; Stress: F_(1, 18)_ = 1.77, *P* = 0.20. **(c)** Trial-averaged Z-scored Δf/F CeA→DMS GCaMP8s fluorescence changes around unpredicted food-pellet reward delivery. **(d)** Trial-averaged quantification CeA→DMS GCaMP8s Z-scored Δf/F AUC during the 3-s period prior to and following reward collection. Stress x Reward: F_(1, 20)_ = 11.79, *P* = 0.02; Reward: F_(1, 20)_ = 8.14, *P* = 0.01; Stress F_(1, 20)_ = 4.49, *P* = 0.05. **(e)** Trial-averaged Z-scored Δf/F BLA→DMS GCaMP8s fluorescence changes around unpredicted footshock. **(f)** Trial-averaged quantification of BLA→DMS GCaMP8s Z-scored Δf/F AUC during the 1-s acute shock response compared to a 1-s pre-shock baseline. Shock: F_(1, 18)_ = 8.533, *P* = 0.01; Stress: F_(1, 18)_ = 0.1433, *P* = 0.71; Stress x Shock F_(1, 18)_ = 1.725, *P* = 0.21 **(g)** Trial-averaged quantification of BLA→DMS GCaMP8s Z-scored Δf/F AUC during 2-s post-shock period. t_(18)_ = 2.26, *P* = 0.04. **(h)** Trial-averaged Z-scored Δf/F CeA→DMS GCaMP8s fluorescence changes around unpredicted footshock. **(i)** Trial-averaged quantification of CeA→DMS GCaMP8s Z-scored Δf/F AUC during the 1-s acute shock response, compared to baseline. Shock: F_(1, 20)_ = 28.24, *P* < 0.0001; Stress: F_(1, 20)_ = 0.22, *P* = 0.64; Stress x Shock: F_(1, 20)_ = 3.201, *P* = 0.09. **(j)** Trial-averaged quantification of CeA→DMS GCaMP8s Z-scored Δf/F AUC during 2-s post-shock period. t_(20)_ = 0.8798, *P* = 0.39. BLA Control N = 8 (4 male), BLA Stress N = 12 (5 male). CeA Control N = 11 (6 male), CeA Stress N = 11 (4 male). Males = solid lines, Females = dashed lines. BLA→DMS projections are activated by unpredicted rewards and this is attenuated by prior chronic stress. Conversely, CeA→DMS projections are not normally robustly activated by unpredicted rewards, but are activated by unpredicted rewards following chronic stress. Interestingly, unpredicted rewards robustly activated CeA→DMS projections here, but rewards did not evoke such a response early in instrumental training (Figure 2m). Rather rewards responses developed with training. This indicates that stress-induced engagement of the CeA→DMS pathway may require repeated reward experience, which may reflect engagement of this pathway with repeated reinforcement and/or opportunity to learn the value or salience of the reward. We speculate this CeA→DMS engagement could be a compensatory mechanism triggered in response to the lack of engagement of the BLA→DMS pathway. Both BLA→DMS and CeA→DMS pathways are acutely activated by unpredicted footshock regardless of prior stress. Though chronic stress reduces post-shock activity in the BLA→DMS pathway.

**Extended Data Figure 2-5:**
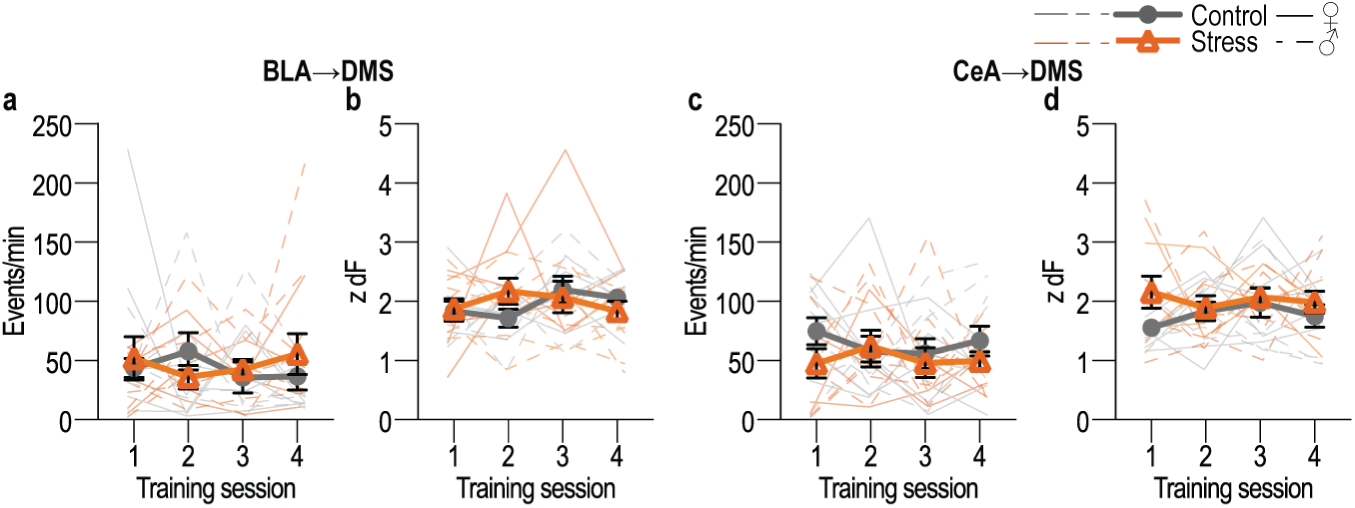
Chronic stress does not affect spontaneous calcium activity in BLA→DMS or CeA→DMS projections. **(a-b)** Frequency (a; Training: F_(2.41, 45.69)_ = 0.17, *P* = 0.88; Stress: F_(1, 19)_ = 0.08, *P* = 0.78; Training x Stress: F_(3, 57)_ = 0.85, P = 0.47) and amplitude (b; Training: F_(2.48, 47.10)_ = 0.86, *P* = 0.45; Stress: F_(1, 19)_ = 0.034, *P* = 0.85; Training x Stress: F_(3, 57)_ = 1.37, *P* = 0.26) of Z-scored Δf/F spontaneous calcium activity of BLA→DMS projections during the 3-min baseline period prior to each training session in handled control and stressed mice. **(c-d)** Frequency (c; Training: F_(2.70, 53.97)_ = 0.21, *P* = 0.88; Stress F_(1, 20)_ = 3.03, *P* = 0.10; Training x Stress: F_(3, 60)_ = 0.55, *P* = 0.65) and amplitude (d; Training: F_(2.59, 51.83)_ = 0.32, *P* = 0.78; Stress: F_(1, 20)_ = 3.70, *P* = 0.07; Training x Stress: F_(3, 60)_ = 0.75, *P* = 0.52) of Z-scored Δf/F spontaneous calcium activity of CeA→DMS projections during the 3-min baseline period prior to each training session handled control and stressed mice. Males = solid lines, Females = dashed lines. Chronic stress did not alter baseline spontaneous calcium activity in either pathway.

**Extended Data Figure 3-1:**
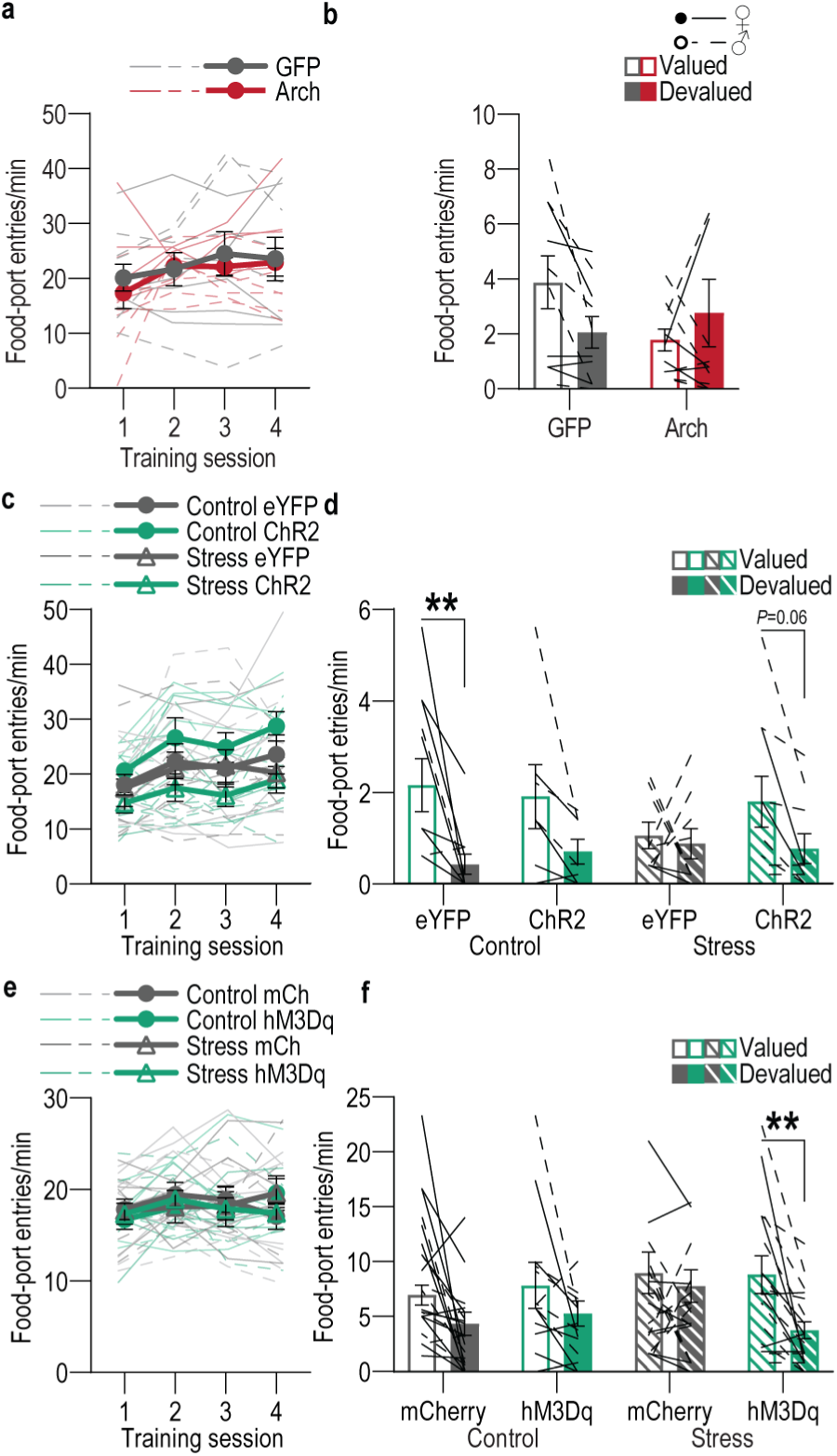
Food-port entries during training with BLA→DMS manipulations and devaluation probe tests. **(a-b)** Optogenetic inactivation of BLA→DMS projections at reward during instrumental learning. **(a)** Food-port entries across training. Training: F_(2.03, 38.55)_ = 3.30, *P* = 0.05; Virus: F_(1, 19)_ = 0.14, *P* = 0.71; Training x Virus: F_(3, 57)_ = 0.43, *P* = 0.73. **(b)** Food-port entry rates during devaluation probe tests. Stress x Value: F_(1, 19)_ = 4.38, *P* = 0.05; Stress: F_(1, 19)_ = 0.47, *P* = 0.50; Value: F_(1, 19)_ = 0.39, *P* = 0.54. eYFP N = 10 (5 males), Arch N = 11 (5 male). **(c-d)** Optogenetic activation of BLA→DMS projections during post-stress instrumental learning. **(c)** Food-port entry rate across training. Training: F_(2.5, 82.82)_ = 6.47, *P* = 0.001; Stress: F_(1, 33)_ = 3.78, *P* = 0.06; Virus: F_(1, 33)_ = 0.02, *P* = 0.89; Training x Stress: F_(3, 99)_ = 0.67, *P* = 0.57; Training x Virus: F_(3, 99)_ = 0.45, *P* = 0.72; Stress x Virus: F_(1, 33)_ = 2.18, *P* = 0.15; Training x Stress x Virus: F_(3, 99)_ = 0.26, *P* = 0.86. **(d)** Food-port entry rate during the devaluation probe tests. Value: F_(1, 33)_ = 15.65, *P* = 0.0004; Stress: F_(1, 33)_ = 0.23, *P* = 0.63; Virus: F_(1, 33)_ = 0.20, *P* = 0.65; Value x Stress: F_(1, 33)_ = 2.75, *P* = 0.11; Value x Virus: F_(1, 33)_ = 0.09, *P* = 0.76; Virus x Stress: F_(1, 33)_ = 0.17, *P* = 0.68; Value x Stress x Virus: F_(1, 33)_ = 1.73, *P* = 0.20. Control, Value: F_(1, 16)_ = 12.42, *P* = 0.003; Virus: F_(1, 16)_ = 0.0007, *P* = 0.98; Value x Virus: F_(1, 16)_ = 0.40, *P* = 0.53. Stress, Value: F_(1, 17)_ = 3.46, *P* = 0.08; Virus: F_(1, 17)_ = 0.45, *P* = 0.51; Value x Virus: F_(1, 17)_ = 1.71, *P* = 0.21. Control eYFP N = 11 (7 male), Control ChR2 N = 7 (4 males), Stress eYFP N = 9 (2 male), Stress ChR2 N = 10 Stress (3 male). **(e-f)** Chemogenetic activation of BLA→DMS projections during post-stress instrumental learning. **(e)** Food-port entry rate across training. Training: F_(2.55, 84.12)_ = 1.64, *P* = 0.19; Stress: F_(1, 33)_ = 0.05, *P* = 0.95; Virus: F_(1, 33)_ = 0.08, *P* = 0.78; Training x Stress: F_(3, 99)_ = 0.16, *P* = 0.92; Training x Virus: F_(3, 99)_ = 0.21, *P* = 0.89; Stress x Virus: F_(1, 33)_ = 0.02, *P* = 0.89; Training x Stress x Virus: F_(3, 99)_ = 3.07, *P* = 0.03. **(f)** Food-port entry rate during the devaluation probe test. Planned comparisons valued v. devalued, Control mCherry: t_(20)_ = 1.88, *P* = 0.07; Control hM3Dq: t_(10)_ = 1.32, *P* = 0.20; Stress mCherry: t_(16)_ = 0.75, *P* = 0.46; Stress hM3Dq: t_(18)_ = 3.36, *P* = 0.002. Control mCherry N = 12 (7 male), Stress mCherry N = 9 (5 male), Stress hM3Dq N = 10 Stress (5 male). Males = solid lines, Females = dashed lines. \*\**P* < 0.01.

**Extended Data Figure 3-2:**
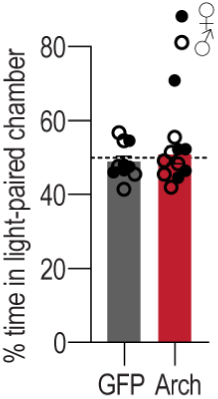
Inhibition of BLA terminals in DMS is not rewarding or aversive. Following training and testing (Figure 3h-n) mice receive a real-time place preference test in which 1 side of a 2-chamber apparatus was paired with optogenetic inhibition of BLA axons and terminals in the DMS. Average percent time spent in light-paired chamber across 2, 10-minute sessions (one with light paired with each side). t_(19)_ = 0.65, *P* = 0.5. eYFP N = 10 (5 male), Arch N = 11 (5 male). Males = closed circles, Females = open circles.

**Extended data Figure 4-1:**
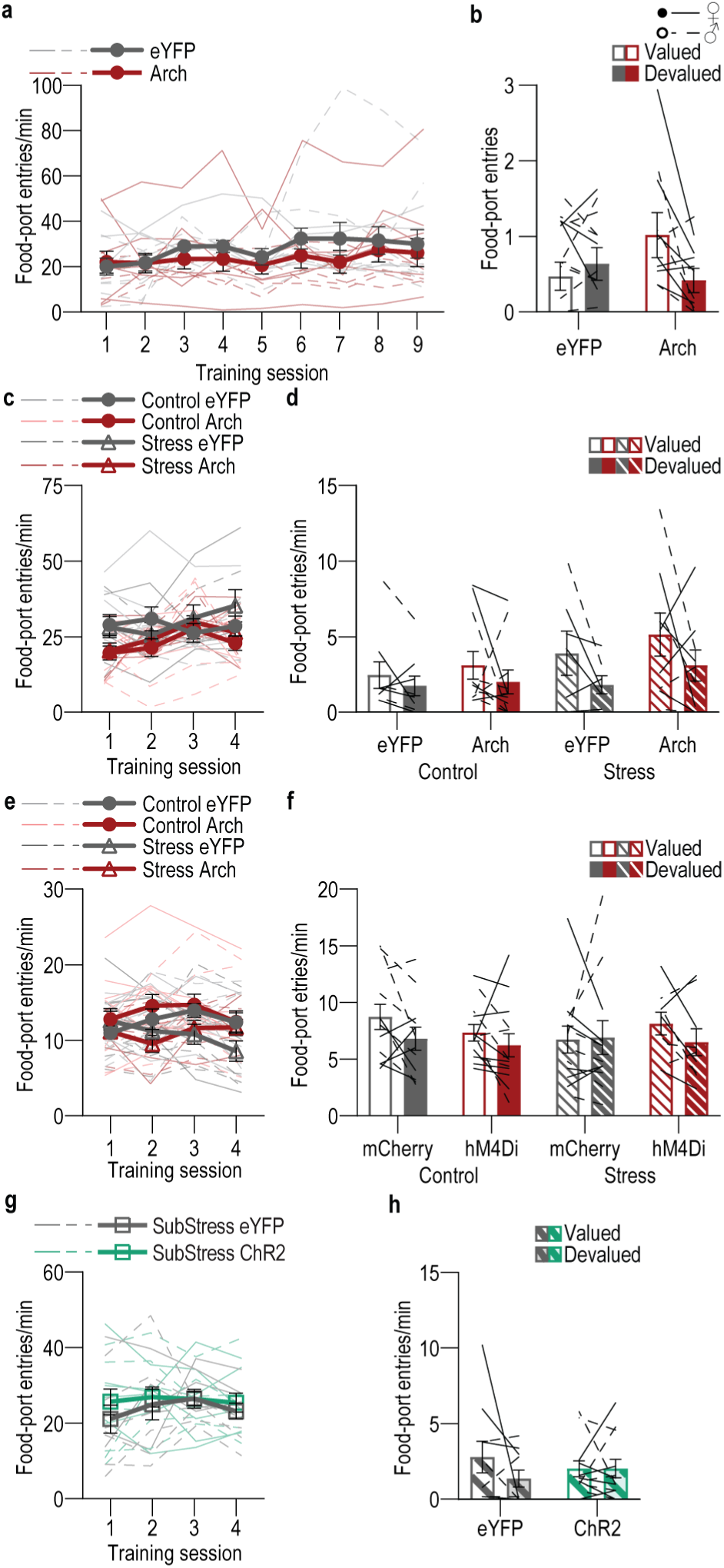
Food-port entries during training with CeA→DMS manipulations and devaluation probe tests. **(a-b)** Optogenetic inhibition of CeA→DMS projections during instrumental overtraining. **(a)** Food-port entry rates across training. Training: F_(2.29, 45.82)_ = 1.81, *P* = 0.17; Virus: F_(1, 20)_ = 0.67, *P* = 0.42; Training x Virus: F_(8, 160)_ = 0.60, *P* = 0.77. **(b)** Food-port entry rates during the devaluation probe tests. Virus x Value: F_(1, 20)_ = 4.51, *P* = 0.046; Value: F_(1, 20)_ = 1.47, *P* = 0.24; Virus: F_(1, 20)_ = 0.41, *P* = 0.53;. eYFP N = 11 (3 male), Arch N = 11 (7 male). **(c-d)** Optogenetic inactivation of CeA→DMS projections at reward during post-stress learning. **(c)** Food-port entry rates across training. Training: F_(2.63, 84.18)_ = 3.21, *P* = 0.03; Stress: F_(1, 32)_ = 0.60, *P* = 0.44; Virus: F_(1, 32)_ = 4.75, *P* = 0.04; Training x Stress: F_(3, 96)_ = 1.55, *P* = 0.21; Training x Virus: F_(3, 96)_ = 2.42, *P* = 0.07; Stress x Virus: F_(1, 32)_ = 0.04, *P* = 0.84; Training x Stress x Virus: F_(3, 96)_ = 1.14, *P* = 0.34. **(k)** Food-port entry rate during the devaluation probe test. Value x Stress x Virus: F_(1, 32)_ = 0.03, *P* = 0.86; Value: F_(1, 32)_ = 6.44, *P* = 0.02; Stress: F_(1, 32)_ = 2.02, *P* = 0.16; Virus: F_(1, 32)_ = 1.09, *P* = 0.30; Value x Stress: F_(1, 3)_ = 0.99, *P* = 0.33; Value x Virus: F_(1, 32)_ = 0.02, *P* = 0.89; Virus x Stress: F_(1, 32)_ = 0.24, *P* = 0.63. *Control groups,* Value x Virus: F_(1, 18)_ = 0.09, *P* = 0.77; Value: F_(1, 18)_ = 1.99, *P* = 0.17; Virus: F_(1, 18)_ = 0.21, *P* = 0.65. *Stress groups,* Value x Virus: F_(1, 14)_ = 0.0005, *P* = 0.98; Value: F_(1, 14)_ = 3.94, *P* = 0.06; Virus: F_(1, 14)_ = 0.85, *P* = 0.87. Control eYFP N = 9 (5 male), Control Arch N = 11 (4 male), Stress eYFP N = 7 (6 male), Stress Arch N = 9 (5 male). **(e-f)** Chemogenetic inhibition of CeA→DMS projections during post-stress instrumental learning. **(e)** Food-port entry rates across training. Training: F_(1.85, 75.67)_ = 2.02, *P* = 0.14; Stress: F_(1, 41)_ = 4.42, *P* = 0.04; Virus: F_(1, 41)_ = 0.41, *P* = 0.53; Training x Stress: F_(3, 123)_ = 3.08, *P* = 0.03; Training x Virus: F_(3, 123)_ = 0.64, *P* = 0.59; Stress x Virus: F_(1, 41)_ = 0.20, *P* = 0.66; Training x Stress x Virus: F_(3, 123)_ = 3.23, *P* = 0.02. **(f)** Food-port entry rates during the devaluation probe tests. Planned comparisons valued v. devalued, Control mCherry: t_(11)_ = 1.94, *P* = 0.06; Control hM3Dq: t_(12)_ = 0.38, *P* = 0.71; Stress mCherry: t_(10)_ = 0.05, *P* = 0.96; Stress hM3Dq: t_(8)_ = 0.47, *P* = 0.64. Control mCherry N = 12 (5 male), Control hM4Di N = 13 (8 male), Stress mCherry N = 11 (5 male), Stress hM4Di N = 9 (4 male). **(g-h)** Optogenetic stimulation of CeA→DMS projections at reward during learning following subthreshold once daily stress (SubStress). **(g)** Food-port entry rate across training. Training: F_(1.73, 34.50)_ = 0.89, *P* = 0.41; Virus: F_(1, 20)_ = 0.46, *P* = 0.51; Training x Virus: F_(3, 60)_ = 0.39, *P* = 0.76. **(g)** Food-port entry rate during the devaluation probe test. Virus x Value: F_(1, 20)_ = 1.37, *P* = 0.26; Virus: F_(1, 20)_ = 0.005, *P* = 0.94; Value: F_(1, 20)_ = 1.36, *P* = 0.26. eYFP N = 10 (4 male), ChR2 N = 12 (6 male). Males = solid lines, Females = dashed lines.

**Extended data Figure 4-2:**
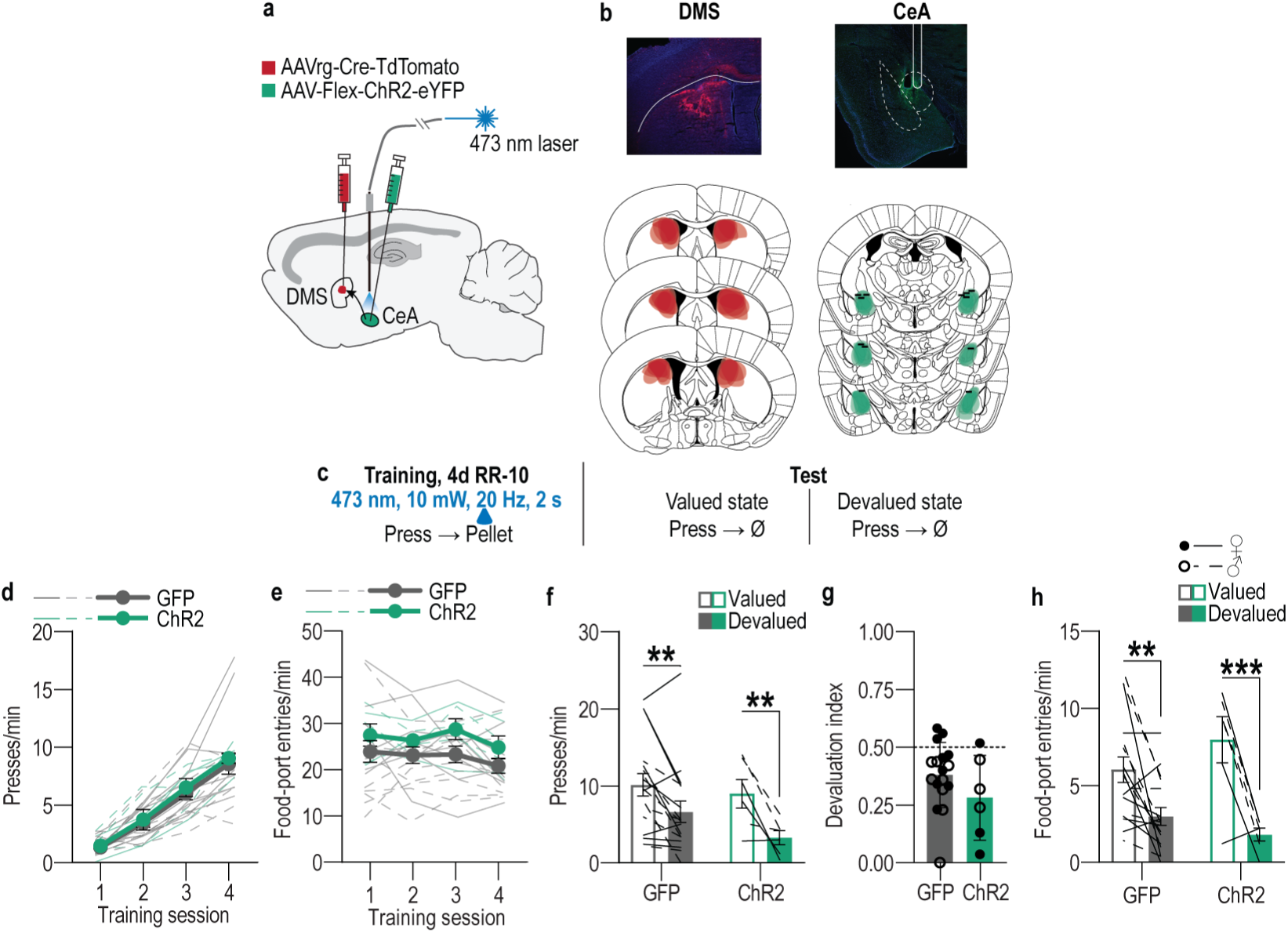
Optogenetic stimulation of CeA→DMS projections in control mice. **(a)** We used an intersectional approach to express the excitatory opsin Channelrhodopsin 2 (ChR2), or a fluorophore control in DMS-projecting CeA neurons and implanted optic fibers above the CeA. **(b)** Representative images of retro-cre expression in DMS and immunofluorescent staining of cre-dependent ChR2 expression in CeA and schematic representation of retro-cre in DMS and cre-dependent ChR2 expression in CeA for all subjects. **(c)** Procedure schematic. Lever presses earned food pellet rewards on a random-ratio (RR) reinforcement schedule. We used blue light (473 nm, 10 mW, 20 Hz, 25-ms pulse width, 2 s) to stimulate CeA→DMS neurons during the collection of each earned reward in mice without a history of stress. Mice were then given a lever-pressing probe test in the Valued state, prefed on untrained food-pellet type to control for general satiety, and Devalued state prefed on trained food-pellet type to induce sensory-specific satiety devaluation (order counterbalanced). **(d)** Press rates across training. Training: F_(1.85, 38.75)_ = 62.18, *P* < 0.0001; Virus: F_(1, 21)_ = 0.23, *P* = 0.64; Training x Virus: F_(3, 63)_ = 0.05, *P* = 0.98. **(e)** Food-port entries across training. Training: F_(2.42, 50.77)_ = 2.00, *P* = 0.14; Virus: F_(1, 21)_ = 1.85, *P* = 0.19; Training x Virus: F_(3, 63)_ = 0.22, *P* = 0.88. **(f)** Press rate during the devaluation probe test. Value: F_(1, 21)_ = 20.32, *P* = 0.0002; Virus: F_(1,21)_ = 0.92, *P* = 0.35; Virus x Value: F_(1, 21)_ = 1.17, *P* = 0.29. **(g)** Devaluation index. t_(21)_ = 1.37, *P* = 0.19. **(h)** Food-port entries during the devaluation probe tests. Value: F_(1, 21)_ = 30.07, *P* < 0.0001; Virus: F_(1, 21)_ = 0.12, *P* = 0.73; Virus x Value: F_(1, 21)_ = 3.45, *P* = 0.08. eYFP N = 17 (9 male), ChR2 N = 6 (3 male). ** *P* < 0.01, *** *P* < 0.001. Optogenetic activation of CeA→DMS projections at reward during learning neither affects affect acquisition of the lever-press behavior, nor the action-outcome learning needed to support flexible goal-directed decision making during the devaluation test.

**Extended data Figure 4-3:**
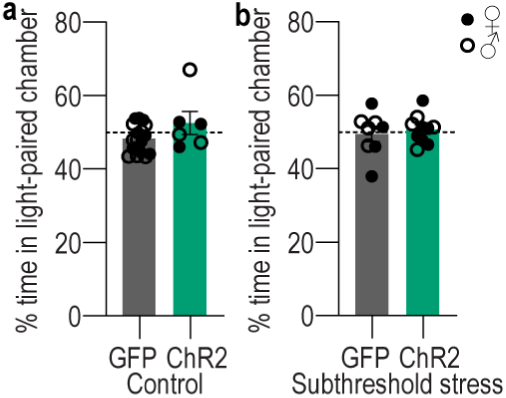
Activation of CeA→DMS projections is neither rewarding or aversive. Following training and testing mice receive a real-time place preference test in which 1 side of a 2-chamber apparatus was paired with optogenetic stimulation of DMS-projecting CeA neurons. **(a)** Average percent time spent in light paired chamber across 2, 10-minute sessions (one with light paired with each side) in handled control subjects. t_(21)_ = 1.75, *P* = 0.10. eYFP N = 17 (9 male), ChR2 N = 6 (3 male). **(b)** Average percent time spent in light paired chamber across 2, 10-minute sessions (one with light paired with each side) in subjects with a prior once/daily stress for 14 d. t_(16)_ = 0.52, *P* = 0.61. eYFP N = 8 (4 male), ChR2 N = 10 (6 male). Males = closed circles, Females = open circles.

## SUPPLEMENTAL TABLES

**Supplemental Table 1:**
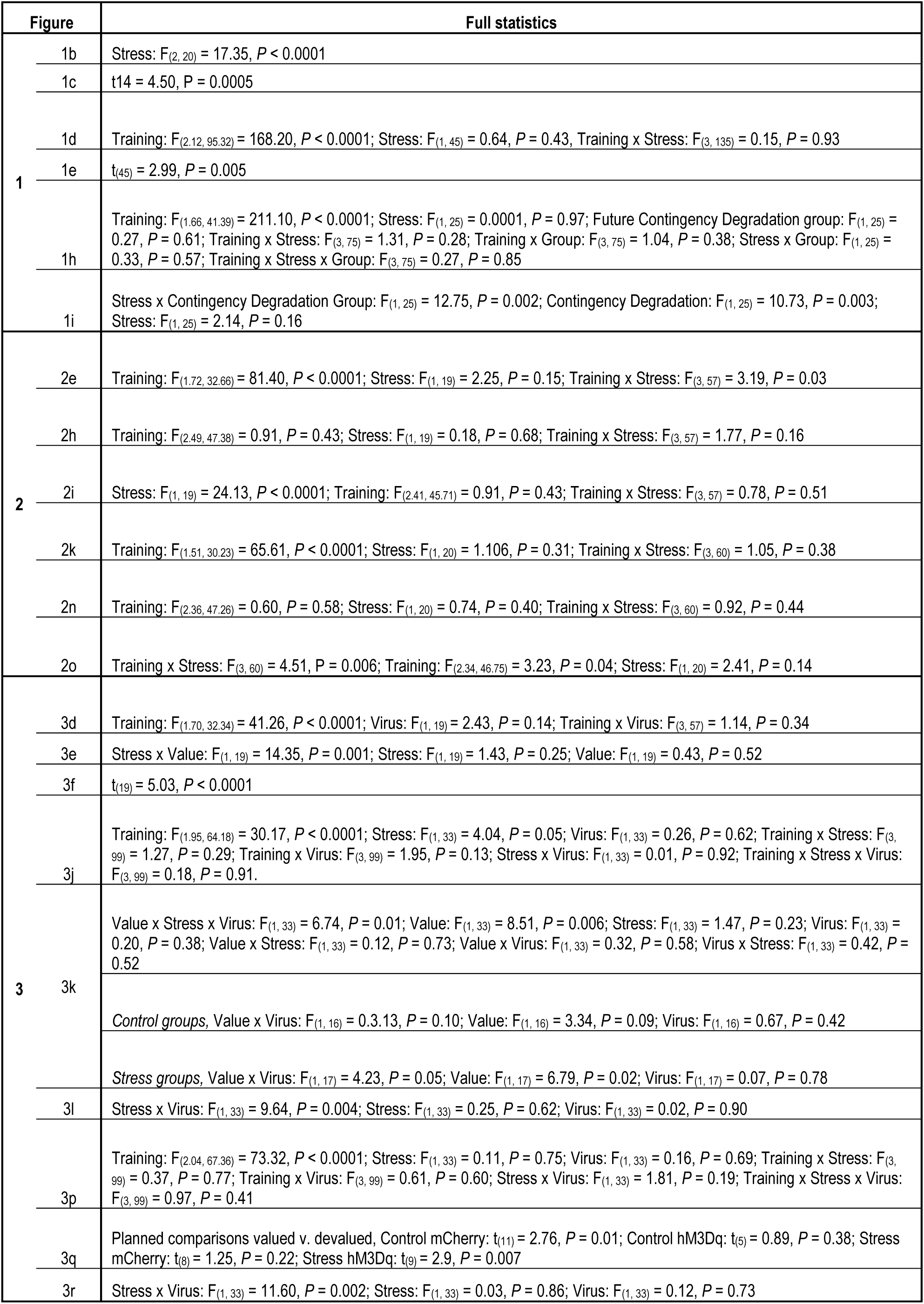
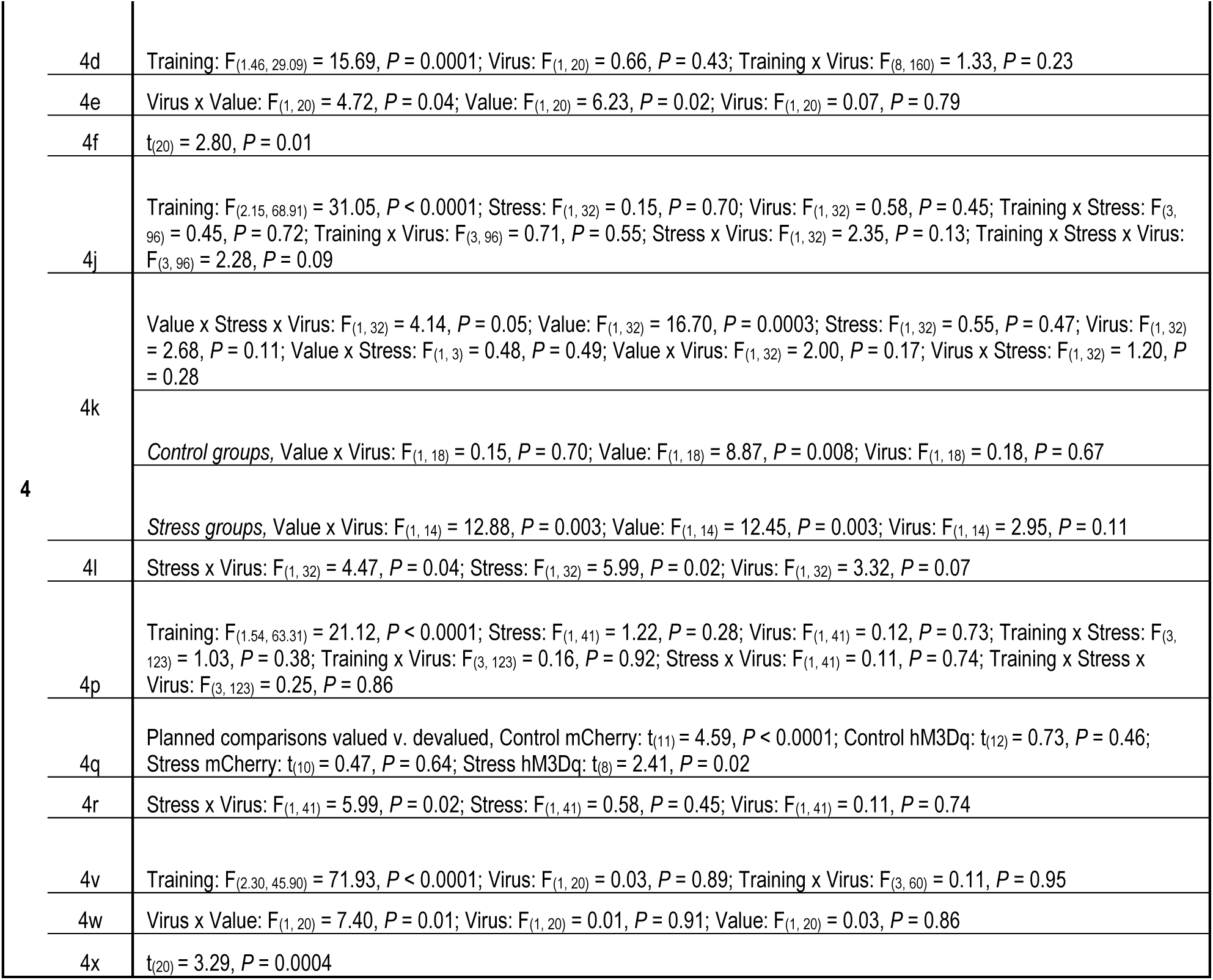
Full statistical reporting for main text data.

**Supplemental Table 2:** Body weight across training. Values reflect average weight in grams ± s.e.m.

| Body Weight |  |  |  |  |
| --- | --- | --- | --- | --- |
| Panel | Group | Training Start | Training End | Statistics |
| Figure 1 d-f | Control | 21.67 ± 0.82 | 21.54 ± 0.90 | Stress x Time: F (1, 45) = 0.08, P = 0.77; Stress: F (1, 45) = 0.60, P = 0.44; Time: F (1, 45) = 1.79, P = 0.19 |
|  | Stress | 20.79 ± 0.81 | 20.60 ± 0.82 |  |
| Figure 1 h-i | Control Non-Deg | 21.51 ± 1.19 | 22.25 ± 1.37 | Training x Stress x Degradation: F (1, 25) = 0.04, P = 0.85; Stress x Degradation: F (1, 25) = 0.65, P = 0.43; Training x Degradation: F (1, 25) = 0.12, P = 0.73; Training x Stress: F (1, 25) = 1.51, P = 0.23; Degradation: F (1, 25) = 0.58, P = 0.45; Stress: F (1, 25) = 2.85, P = 0.10; Training: F (1, 25) = 13.84, P = 0.001 |
|  | Control Deg | 23.48 ± 1.30 | 24.27 ± 1.34 |  |
|  | Stress Non-Deg | 20.60 ± 1.24 | 20.91 ± 1.03 |  |
|  | Stress Deg | 20.46 ± 1.40 | 20.94 ± 1.30 |  |
| Figure 2 | Control CeA | 20.06 ± 0.94 | 20.24 ± 0.71 | Stress x Training: F (1, 20) = 0.16, P = 0.70; Stress: F (1, 20) = 1.02, P = 0.32; Training: F (1, 20) = 0.11, P = 0.75 |
|  | Stress CeA | 19.12 ± 0.60 | 19.10 ± 0.72 |  |
|  | Control BLA | 20.54 ± 1.00 | 20.25 ± 0.92 | Stress x Training: F (1, 19) = 2.67, P = 0.12; Stress: F (1, 19) = 2.24, P = 0.15; Training: F (1, 19) = 0.02, P = 0.88 |
|  | Stress BLA | 18.66 ± 0.60 | 19.01 ± 0.56 |  |
| Figure 3 a-f | eYFP | 21.91 ± 1.30 | 21.25 ± 1.24 | Virus x Training: F (1, 19) = 8.58, P = 0.009; Virus: F (1, 19) = 0.31, P = 0.59; Training: F (1, 19) = 4.96, P = 0.04 |
|  | Arch | 20.73 ± 0.81 | 20.82 ± 0.75 |  |
| Figure 3 g-l | Control eYFP | 23.59 ± 1.13 | 23.04 ± 0.92 | Training x Virus x Stress: F (1, 33) = 1.31, P = 0.26; Virus x Stress: F (1, 33) = 0.0004, P = 0.98; Training x Stress: F (1, 33) = 1.36, P = 0.25; Training x Virus: F (1, 33) = 1.51, P = 0.23; Stress: F (1, 33) = 6.04, P = 0.02; Virus: F (1, 33) = 0.49, P = 0.49; Training: F (1, 33) = 5.13, P = 0.03 |
|  | Stress eYFP | 21.22 ± 0.96 | 20.67 ± 0.78 |  |
|  | Control ChR2 | 22.55 ± 1.25 | 22.75 ± 1.18 |  |
|  | Stress ChR2 | 20.50 ± 0.91 | 19.98 ± 0.62 |  |
| Figure 3 m-r | Control mCherry | 21.48 ± 1.01 | 21.08 ± 0.97 | Training x Virus x Stress: F (1, 33) = 1.63, P = 0.21; Virus x Stress: F (1, 33) = 0.07, P = 0.80; Training x Stress: F (1, 33) = 0.24, P = 0.63; Training x Virus: F (1, 33) = 1.15, P = 0.29; Stress: F (1, 33) = 0.17, P = 0.68; Virus: F (1, 33) = 0.76, P = 0.39; Training: F (1, 33) = 0.16, P = 0.69 |
|  | Stress mCherry | 20.65 ± 0.86 | 20.53 ± 0.97 |  |
|  | Control hM3Gq | 19.91 ± 1.34 | 20.35 ± 1.64 |  |
|  | Stress hM3Gq | 20.06 ± 0.75 | 19.87 ± 0.72 |  |
| Figure 4 a-f | eYFP | 21.59 ± 0.96 | 22.10 ± 1.02 | Virus x Training: F (1, 20) = 0.03, P = 0.87; Training: F (1, 20) = 0.35, P = 0.56; Virus: F (1, 20) = 0.09, P = 0.76 |
|  | Arch | 21.6 ± 0.95 | 22.50 ± 0.88 |  |
| Figure 4 g-l | Control mCherry | 22.66 ± 1.19 | 22.26 ± 0.98 | Training x Virus x Stress: F (1, 32) = 0.25, P = 0.62; Virus x Stress: F (1, 32) = 0.14, P = 0.71; Training x Stress: F (1, 32) = 2.15, P = 0.15; Training x Virus: F (1, 32) = 0.46, P = 0.50; Stress: F (1, 32) = 0.09, P = 0.76; Virus: F (1, 32) = 4.76, P = 0.04; Training: F (1, 32) = 6.85, P = 0.01 |
|  | Stress mCherry | 22.76 ± 0.99 | 22.03 ± 0.82 |  |
|  | Control Arch | 20.03 ± 0.86 | 20.04 ± 0.85 |  |
|  | Stress Arch | 21.00 ± 0.92 | 20.36 ± 0.92 |  |
| Figure 4 m-r | Control mCherry | 21.81 ± 1.11 | 21.94 ± 1.11 | Training x Virus x Stress: F (1, 41) = 0.61, P = 0.44; Virus x Stress: F (1, 41) = 0.31, P = 0.58; Training x Stress: F (1, 41) = 0.01, P = 0.94; Training x Virus: F (1, 41) = 0.39, P = 0.54; Stress: F (1, 41) = 0.96, P = 0.33; Virus: F (1, 41) = 0.18, P = 0.67; Training: F (1, 41) = 1.11, P = 0.30 |
|  | Stress mCherry | 21.24 ± 1.03 | 21.66 ± 0.97 |  |
|  | Control hM4Gi | 22.77 ± 0.90 | 22.96 ± 0.86 |  |
|  | Stress hM4Gi | 21.34 ± 1.08 | 21.30 ± 0.98 |  |
| Figure 4-3 | Control GFP | 20.87 ± 0.75 | 20.81 ± 0.70 | Virus x Training: F (1, 21) = 0.79, P = 0.38; Virus: F (1, 21) = 0.94, P = 0.34; Training: F (1, 21) = 0.15, P = 0.70 |
|  | Control ChR2 | 19.34 ± 1.39 | 19.50 ± 1.4 |  |
| Figure 4 s-x | Sub-Stress GFP | 20.33 ± 0.89 | 20.14 ± 0.69 | Virus x Training: F (1, 20) = 0.03, P = 0.87; Virus: F (1, 20) = 0.75, P = 0.40; Training: F (1, 20) = 1.78, P = 0.20 |
|  | Sub-Stress ChR2 | 21.37 ± 0.86 | 21.13 ± 0.84 |  |

**Supplemental Table 3:** Sensory-specific satiety prefeed consumption. Values reflect average amount consumed in grams ± s.e.m.

|  |  | Pre-feed Consumption |  |  |
| --- | --- | --- | --- | --- |
|  |  | Valued Outcome | Devalued Outcome | Statistics |
| Figure 1 d-f | Control | 1.33 ± 0.09 | 1.32 ± 0.10 | Stress x Value: F (1, 45) = 0.18, P = 0.67; Stress: F (1, 45) = 0.13, P = 0.72; Value: F (1, 45) = 0.33, P = 0.57 |
|  | Stress | 1.33 ± 0.08 | 1.25 ± 0.09 |  |
| Figure 3 a-f | GFP | 1.57 ± 0.19 | 1.40 ± 0.07 | Virus x Value: F (1, 19) = 0.46, P = 0.50; Virus: F (1, 19) = 0.007, P = 0.94; Value: F (1, 19) = 0.21, P = 0.65 |
|  | Arch | 1.48 ± 0.19 | 1.51 ± 0.16 |  |
| Figure 3 g-l | Control eYFP | 1.48 ± 0.12 | 1.35 ± 0.10 | Value x Virus x Stress: F (1, 33) = 0.20, P = 0.66; Virus x Stress: F (1, 33) = 0.36, P = 0.55; Value x Stress: F (1, 33) = 1.94, P = 0.90; Value x Virus: F (1, 33) = 0.02, P = 0.90; Stress: F (1, 33) = 11.20, P = 0.002; Virus: F (1, 33) = 5.48, P = 0.03; Value: F (1, 33) = 0.37, P = 0.55 |
|  | Stress eYFP | 1.07 ± 0.08 | 1.30 ± 0.20 |  |
|  | Control ChR2 | 1.66 ± 0.16 | 1.64 ± 0.11 |  |
|  | Stress ChR2 | 1.25 ± 0.16 | 1.41 ± 0.14 |  |
| Figure 3 m-r | Control mCherry | 1.10 ± 0.68 | 1.39 ± 0.14 | Value x Virus: F (1, 33) = 0.09, P = 0.76; Virus x Stress: F (1, 33) = 0.70, P = 0.41; Value x Stress: F (1, 33) = 7.30, P = 0.01; Value x Virus: F (1, 33) = 0.02, P = 0.90; Stress: F (1, 33) = 1.44, P = 0.24; Virus: F (1, 33) = 0.42, P = 0.52; Value: F (1, 33) = 0.01, P = 0.92 |
|  | Stress mCherry | 1.56 ± 0.13 | 1.28 ± 0.09 |  |
|  | Control hM3Gq | 1.27 ± 0.15 | 1.47 ± 0.17 |  |
|  | Stress hM3Gq | 1.53 ± 0.10 | 1.28 ± 0.14 |  |
| Figure 4 a-f | eYFP | 1.33 ± 0.11 | 1.57 ± 0.17 | Value x Virus: F (1, 20) = 1.62, P = 0.22; Value: F (1, 20) = 1.20, P = 0.29; Virus: F (1, 20) = 0.20, P = 0.66 |
|  | Arch | 1.35 ± 0.13 | 1.24 ± 0.13 |  |
| Figure 4 g-l | Control eYFP | 1.62 ± 0.16 | 1.68 ± 0.12 | Value x Virus x Stress: F (1, 32) = 1.07, P = 0.31, Virus x Stress: F (1, 32) = 0.01, P = 0.94; Value x Stress: F (1, 32) = 0.10, P = 0.75; Value x Virus: F (1, 32) = 0.31, P = 0.58; Stress: F (1, 32) = 0.46, P = 0.50; Virus: F (1, 32) = 5.64, P = 0.02; Value: F (1, 32) = 0.61, P = 0.44 |
|  | Stress eYFP | 1.50 ± 0.14 | 1.70 ± 0.17 |  |
|  | Control Arch | 1.40 ± 0.13 | 1.52 ± 0.13 |  |
|  | Stress Arch | 1.42 ± 0.16 | 1.31 ± 0.10 |  |
| Figure 4 m-r | Control mCherry | 0.72 ± 0.13 | 0.80 ± 0.13 | Value x Virus x Stress: F (1, 41) = 0.01, P = 0.91; Virus x Stress: F (1, 41) = 0.01, P = 0.94; Value x Stress: F (1, 41) = 0.93, P = 0.34; Value x Virus: F (1, 41) = 0.64, P = 0.43; Stress: F (1, 41) = 2.83, P = 0.10; Virus: F (1, 41) = 0.14, P = 0.71; Value: F (1, 41) = 0.19, P = 0.66 |
|  | Stress mCherry | 0.57 ± 0.16 | 0.54 ± 0.12 |  |
|  | Control hM4Gi | 0.69 ± 0.07 | 0.53 ± 0.13 |  |
|  | Stress hM4Gi | 0.80 ± 0.14 | 0.79 ± 0.14 |  |
| Figure 4-3 | Control GFP | 1.31 ± 0.09 | 1.47 ± 0.12 | Virus x Value: F (1, 25) = 0.016, P = 0.90; Virus: F (1, 25) = 0.24, P = 0.63; Value: F (1, 25) = 1.96, P = 0.17 |
|  | Control ChR2 | 1.36 ± 0.16 | 1.55 ± 0.15 |  |
| Figure 4 s-x | Sub-Stress GFP | 1.49 ± 0.18 | 1.54 ± 0.133 | Virus x Value: F (1, 20) = 0.04, P = 0.84; Virus: F (1, 20) = 1.11, P = 0.30; Value: F (1, 20) = 0.01, P = 0.92 |
|  | Sub-Stress ChR2 | 1.66 ± 0.13 | 1.65 ± 0.14 |  |

**Supplemental Table 4:** Average post-probe-test choice consumption. Values reflect average amount consumed in grams ± s.e.m.

|  |  | Average Choice Consumption |  |  |
| --- | --- | --- | --- | --- |
|  |  | Valued Outcome | Devalued Outcome | Statistics |
| Figure 1 d-f | Control | 0.25 ± 0.03 | 0.01 ± 0.01 | Stress x Value: F (1, 26) = 0.002, P=0.97; Stress: F (1, 26) = 0.006, P=0.94; Value: F (1, 26) = 132.10, P<0.0001 |
|  | Stress | 0.25 ± 0.03 | 0.01 ± 0.01 |  |
| Figure 3 a-f | GFP | 0.25 0.03 | 0.02 0.01 | Virus x Value: F (1, 19) = 0.14, P=0.71; Virus: F (1, 19) = 1.21, P=0.29; Value: F (1, 19) = 118.00, P<0.0001 |
|  | Arch | 0.30 0.04 | 0.05 0.02 |  |
| Figure 3 g-l | Control eYFP | 0.31 ± 0.07 | 0.05 ± 0.03 | Value x Virus x Stress: F (1, 33) = 0.05, P = 0.82; Virus x Stress: F (1, 33) = 0.09, P = 0.77; Value x Stress: F (1, 33) = 0.46, P = 0.50; Value x Virus: F (1, 33) = 1.16, P = 0.29; Stress: F (1, 33) = 1.36, P = 0.25; Virus: F (1, 33) = 0.12, P = 0.73; Value: F (1, 33) = 67.81, P<0.0001 |
|  | Stress eYFP | 0.25 ± 0.06 | 0.02 ± 0.01 |  |
|  | Control ChR2 | 0.36 ± 0.05 | 0.01 ± 0.01 |  |
|  | Stress ChR2 | 0.30 ± 0.05 | 0.01 ± 0.01 |  |
| Figure 3 m-r | Control mCherry | 0.23 ± 0.01 | 0.02 ± 0.01 | Value x Virus x Stress: F (1, 33) = 0.22, P = 0.65; Virus x Stress: F (1, 33) = 5.00, P = 0.03; Value x Stress: F (1, 33) = 0.288, P = 0.10; Value x Virus: F (1, 33) = 1.70, P = 0.20; Stress: F (1, 33) = 0.68, P = 0.42; Virus: F (1, 33) = 0.19, P = 0.67; Value: F (1, 33) = 260.4, P<0.0001 |
|  | Stress mCherry | 0.29 ± 0.03 | 0.03 ± 0.01 |  |
|  | Control hM3Gq | 0.24 ± 0.02 | 0.05 ± 0.03 |  |
|  | Stress hM3Gq | 0.24 ± 0.02 | 0.02 ± 0.01 |  |
| Figure 4 a-f | Control eYFP | 0.24 ± 0.02 | 0.01 ± 0.01 | Virus x Value: F (1, 20) = 0.09, P = 0.77; Virus: F (1, 20) = 0.04, P = 0.85; Value: F (1, 20) = 158.10, P<0.0001 |
|  | Control Arch | 0.25 ± 0.03 | 0.01 ± 0.01 |  |
| Figure 4 g-l | Control eYFP | 0.23 ± 0.04 | 0.02 ± 0.01 | Value x Virus x Stress: F (1, 32) = 0.03, P = 0.86; Virus x Stress: F (1, 32) = 0.01, P>0.99; Value x Stress: F (1, 32) = 0.18, P = 0.67; Value x Virus: F (1, 32) = 2.23, P = 0.15; Stress: F (1, 32) = 0.13, P = 0.72; Virus: F (1, 32) = 2.43, P = 0.13; Value: F (1, 32) = 87.94, P<0.0001 |
|  | Stress eYFP | 0.21 ± 0.03 | 0.02 ± 0.01 |  |
|  | Control Arch | 0.17 ± 0.04 | 0.02 ± 0.01 |  |
|  | Stress Arch | 0.16 ± 0.02 | 0.02 ± 0.01 |  |
| Figure 4 m-r | Control mCherry | 0.16 ± 0.04 | 0.11 ± 0.04 | Value x Virus x Stress: F (1, 41) = 0.01, P = 0.97; Virus x Stress: F (1, 41) = 0.14, P = 0.71; Value x Stress: F (1, 41) = 0.79, P = 0.38; Value x Virus: F (1, 41) = 0.09, P = 0.77; Stress: F (1, 41) = 0.07, P = 0.80; Virus: F (1, 41) = 1.40, P = 0.24; Value: F (1, 41) = 3.78, P = 0.05 |
|  | Stress mCherry | 0.12 ± 0.04 | 0.10 ± 0.04 |  |
|  | Control hM4Gi | 0.19 ± 0.05 | 0.16 ± 0.06 |  |
|  | Stress hM4Gi | 0.21 ± 0.06 | 0.14 ± 0.04 |  |
| Figure 4-3 | Control GFP | 0.27 ± 0.02 | 0.02 ± 0.01 | Virus x Value: F (1, 25) = 0.001, P=0.97; Virus: F (1, 25) = 0.22, P=0.64; Value: F (1, 25) = 160.50, P<0.0001 |
|  | Control ChR2 | 0.28 ± 0.03 | 0.02 ± 0.01 |  |
| Figure 4 s-x | Sub-Stress GFP | 0.26 ± 0.03 | 0.02 ± 0.01 | Virus x Value: F (1, 20) = 0.55, P = 0.47; Virus: F (1, 20) = 1.00, P = 0.33; Value: F (1, 20) = 78.75, P<0.0001 |
|  | Sub-Stress ChR2 | 0.22 ± 0.04 | 0.02 ± 0.01 |  |

**Supplemental Table 5:** Key reagents information.

| Category | Item | Vendor | Catalog # | Lot # | Titer |
| --- | --- | --- | --- | --- | --- |
| ELISA Kit | Highly sensitive corticosterone ELISA kit | Enzo Life Sciences | ADI-900-097 | 02252009C |  |
| Viruses/Tracers | AAV8-hSyn-mCherry | UNC Vector Core | 114472 | v113707 | 2.6 x 10<sup>13</sup> gc/mL |
|  | Fluorogold | Santa Cruz Biotechnology | sc-358883 | A1223 |  |
|  | AAV9-Syn-FLEX-jGCaMP8s-WPRE | Addgene | 162377 | v118930 | 5 x 10<sup>12</sup> gc/mL |
|  | AAVrg-hSyn-Cre-P2A-tdTomato | Addgene | 107738 | v75881 | 1.5 x 10<sup>13</sup> pp/mL |
|  | AAV2-hSyn-DIO-hM3D(Gq)-mCherry | Addgene | 44361 | v97910 | 2.0 x 10<sup>13</sup> gc/mL |
|  | AAV2-hSyn-DIO-hM4D(Gi)-mCherry | Addgene | 44362 | v68359 | 1.5 x 10<sup>13</sup> gc/mL |
|  | AAV2-hSyn-DIO-mCherry | Addgene | 50459 | v107704 | 1.6 x 10<sup>13</sup> gc/mL |
|  | AAV2-hSyn-DIO-mCherry | Addgene | 50459 | v54505 | 1.8 x 10<sup>13</sup> gc/mL |
|  | AAV2-hSyn-DIO-mCherry | Addgene | 50459 | v122065 | 2.1 x 10<sup>13</sup> gc/mL |
|  | AAV8-hSyn-eYFP | Addgene | 50465 | v53294 | 2.5 x 10<sup>13</sup> pp/mL |
|  | AAV8-hSyn-DIO-GFP | Stanford Vector Core | GVVC-AAV-206 | 7463 | 1 x 10<sup>13</sup> vg/mL |
|  | AAV8-hSyn-DIO-ChR2-eYFP | Stanford Vector Core | GVVC-AAV-207 | 7464 | 7.5 x 10<sup>12</sup> vg/mL |
|  | AAV-DJ-hSyn-eYFP | Stanford Vector Core | GVVC-AAV-16 | 6708 | 2.5 x 10<sup>13</sup> gc/mL |
|  | AAV-DJ-hSyn-eArch-eYFP | Stanford Vector Core | GVVC-AAV-146 | 4799 | 1 x 10<sup>13</sup> gc/mL |
|  | AAV-DJ-hSyn-eArch-eYFP | Stanford Vector Core | GVVC-AAV-146 | 5105 | 9.3 x 10<sup>12</sup> gc/mL |
|  | CAV2-Cre-GFP | Plateforme de Vectorologie de Montpellier | N/A | N/A | 1.42 x 10<sup>13</sup> pp/mL |
|  | CAV2-Cre-GFP | Plateforme de Vectorologie de Montpellier | N/A | N/A | 9.6 x 10<sup>12</sup> pp/mL |
|  | AAV8-hSyn-FLEX-TVA-P2A-GFP-2A-oG | Salk Vector Core | 85225 | N/A | 2.53 x 10<sup>12</sup> gc/mL |
|  | AAV8-hSyn-FLEX-TVA-P2A-GFP-2A-oG | Salk Vector Core | 85225 | N/A | 1.87 x 10<sup>12</sup> gc/mL |
|  | EnvA-Gdeleted-Rabies-mCherry | Salk Vector Core | 32636 | N/A | 1.0 x 10<sup>8</sup> TU/mL |
| Antibodies | Chicken polyclonal anti-mCherry primary antibody | Abcam | ab205402 | GR3368071-2, GR3271744-8 |  |
|  | Chicken polyclonal anti-GFP primary antibody | Abcam | ab13970 | 1018753-7, 1018753-5, 1018753-11, 1018753-12 |  |
|  | Living Colors Rabbit DsRed Polyclonal Antibody | Takara Bio | 632496 | 1904182, 2210019 |  |
|  | Goat anti-Rabbit IgG Secondary Antibody, Alexa Fluor 594 | Invitrogen | A11012 | 2433881, 2119134 |  |
|  | Goat anti-Chicken IgY Secondary Antibody, Alexa Fluor 488 | Abcam | ab150169 | GR3437715-3 |  |

## Notes

### Competing Interest Statement

The authors have declared no competing interest.

### Summary of Updates

Changes include new experiments, analyses, figures, and text edits. In particular, we have included 3 new experiments. We used contingency degradation to provide evidence that stress promotes habit formation by disrupting the action-outcome knowledge needed for goal-directed decision making (Fig. 1g-i). We used optogenetics to provide evidence that stimulating BLA->DMS projections at the time of earned reward during learning rescues action-outcome learning and prevents premature habit formation in stressed subjects (Fig. 3g-l). We used optogenetics to provide evidence that the CeA->DMS pathway is necessary for the natural habit formation that occurs for routine behaviors. This demonstrates that stress does, indeed, recruit the mechanisms required for normal habit formation (Fig. 4a-f).We have added N to increase the power and rigor of each optogenetic experiment. We now show full 2 x 2 (stress x manipulation) datasets in the main figures.

